## Supplementary File S1 for "Polygenic selection to a changing optimum under self–fertilisation": S1_File.pdf

#### S1 File: Two-locus model of selection on a quantitative trait under partial selfing

Matthew Hartfield and Sylvain Glémin

*In this notebook we analyse the two-locus model*

#### Recursions for two quantitative alleles with selfing and mutation-selection balance

This notebook derives the overall recursions for two quantitative loci assuming the following lifecycle: Selection; reproduction.
There are two alleles per locus: A/a and B/b.
aa, bb genotypes have contributions 0, 0.
AA, BB genotypes have contributions $2\gamma A$, $2\gamma B$.
Effects are additive.
So population phenotypic range goes from 0 to $2(\gamma A + \gamma B)$.

I consider change of the four following haplotypes:
x1 = Frequency of ab
x2 = Frequency of Ab
x3 = Frequency of aB
x4 = Frequency of AB

These are packaged up in genotypes, which are denoted as:
g11: Freq ab/ab
g12: Freq ab/Ab
g13: Freq ab/aB
g14: Freq ab/AB
g22: Freq Ab/Ab
g23: Freq Ab/aB
g24: Freq Ab/AB
g33: Freq aB/aB
g34: Freq aB/AB
g44: Freq AB/AB

#### Converting genotypes to allele frequencies

To simplify downstream analyses, I will convert genotype frequencies to allele frequencies after each recursion stage. The approximation I will use will be to consider the potential IBD state of *haplotypes.* That is:
gii = $xi^2$ + F xi (1-xi) for i ∈ {1,..,4}
gij = 2xi xj (1-F) for i ≠ j

Below is first code for converting haplotype freqs to allele freqs:

In[∘]:=
```
PDtoX =
  {x1 → 1 + pA (-1 + pB) - pB + δ, x2 → pA - pA pB - δ, x3 → pB - pA pB - δ, x4 → pA pB + δ};
```

Then converting genotypes to haplotypes:

In[∘]:=
```
Gtype2Htype = {
    g11 → x1^2 + F x1 (1 - x1),
    g12 → 2 x1 x2 (1 - F),
    g13 → 2 x1 x3 (1 - F),
    g14 → 2 x1 x4 (1 - F),
    g22 → x2^2 + F x2 (1 - x2),
    g23 → 2 x2 x3 (1 - F),
    g24 → 2 x2 x4 (1 - F),
    g33 → x3^2 + F x3 (1 - x3),
    g34 → 2 x3 x4 (1 - F),
    g44 → x4^2 + F x4 (1 - x4)
  };
```

#### Phenotype definitions

In[∘]:=
```
(* Individual phenotypes *)
z11 = 0 - z0;
z12 = αA - z0;
z13 = αB - z0;
z14 = αA + αB - z0;
z22 = 2 * αA - z0;
z23 = αA + αB - z0;
z24 = 2 * αA + αB - z0;
z33 = 2 * αB - z0;
z34 = αA + 2 * αB - z0;
z44 = 2 * αA + 2 * αB - z0;
```

#### Fitness definitions

In[∘]:=
```
(* Stabilising selection function, z0 = 0 *)
W[z_] := Exp[-z^2/(2 Vs)]
```

```
In[]:= (* Individual fitness definitions *)
W11 = W[z11];
W12 = W[z12];
W13 = W[z13];
W14 = W[z14];
W22 = W[z22];
W23 = W[z23];
W24 = W[z24];
W33 = W[z33];
W34 = W[z34];
W44 = W[z44];
(* Mean fitness *)
Wm = g11 * W11 + g12 * W12 + g13 * W13 + g14 * W14 +
    g22 * W22 + g23 * W23 + g24 * W24 + g33 * W33 + g34 * W34 + g44 * W44;
```

```
In[]:= (*γa, γb substitutions if wanting to try small selection approximations *)
(*Note that with notation,
Taylor expansion must be done at the second order*)
Ssub = {αA → Sqrt[Vs] ξ γA, αB → Sqrt[Vs] γB ξ, z0 → Sqrt[Vs] d0 ξ};
```

#### Recursions following selection

In[]:=

$$g11s = \frac{g11 * W11}{Wm};$$
$$g12s = \frac{g12 * W12}{Wm};$$
$$g13s = \frac{g13 * W13}{Wm};$$
$$g14s = \frac{g14 * W14}{Wm};$$
$$g22s = \frac{g22 * W22}{Wm};$$
$$g23s = \frac{g23 * W23}{Wm};$$
$$g24s = \frac{g24 * W24}{Wm};$$
$$g33s = \frac{g33 * W33}{Wm};$$
$$g34s = \frac{g34 * W34}{Wm};$$
$$g44s = \frac{g44 * W44}{Wm};$$

Checking that all post-selection frequencies sum to 1 (Note: I ran these checks within each section, WITHOUT running previous recursion parts. This way, I checked that the total genotype frequency is conserved solely within each section, in order to save time).

In[ ]:=
```
g11s + g12s + g13s + g14s + g22s + g23s + g24s + g33s + g34s + g44s /.
  (Wm → g11 * W11 + g12 * W12 + g13 * W13 + g14 * W14 + g22 * W22 +
     g23 * W23 + g24 * W24 + g33 * W33 + g34 * W34 + g44 * W44) // FullSimplify
```
Out[ ]=
```
1
```

In the following approximation we assume that pA < pB << 1.
By construction, $\delta \in$ [-pA pB; pA - pA pB]. Assuming that pA, pB are o($\zeta$) then, at least when negative, $\delta$ is $o(\zeta^2)$

#### Recursions following reproduction

Baseline change in haplotypes following complete outcrossing:

In[ ]:=
$$x1S0 = g11s + \frac{(g12s + g13s + g14s)}{2} - r\frac{(g14s - g23s)}{2};$$
$$x2S0 = g22s + \frac{(g12s + g23s + g24s)}{2} + r\frac{(g14s - g23s)}{2};$$
$$x3S0 = g33s + \frac{(g13s + g23s + g34s)}{2} + r\frac{(g14s - g23s)}{2};$$
$$x4S0 = g44s + \frac{(g14s + g24s + g34s)}{2} - r\frac{(g14s - g23s)}{2};$$

These haplotype forms can be compared with selfing equations to determine the change in frequency given a certain proportion of selfing (Hedrick 1980):"

In[ ]:=
$$g11S = S\left(g11s + \frac{\left(g12s + g13s + r^2\ g23s + (1-r)^2\ g14s\right)}{4}\right) + (1-S) * x1S0^2;$$
$$g22S = S\left(g22s + \frac{\left(g12s + g24s + r^2\ g14s + (1-r)^2\ g23s\right)}{4}\right) + (1-S) * x2S0^2;$$
$$g33S = S\left(g33s + \frac{\left(g13s + g34s + r^2\ g14s + (1-r)^2\ g23s\right)}{4}\right) + (1-S) * x3S0^2;$$
$$g44S = S\left(g44s + \frac{\left(g24s + g34s + r^2\ g23s + (1-r)^2\ g14s\right)}{4}\right) + (1-S) * x4S0^2;$$
$$g12S = S\left(\frac{(g12s + r\ (1-r)\ (g14s + g23s))}{2}\right) + (1-S) * 2 * x1S0 * x2S0;$$
$$g13S = S\left(\frac{(g13s + r\ (1-r)\ (g14s + g23s))}{2}\right) + (1-S) * 2 * x1S0 * x3S0;$$
$$g14S = S\left(\frac{(1-r)^2\ g14s + r^2\ g23s}{2}\right) + (1-S) * 2 * x1S0 * x4S0;$$
$$g23S = S\left(\frac{(1-r)^2\ g23s + r^2\ g14s}{2}\right) + (1-S) * 2 * x2S0 * x3S0;$$
$$g24S = S\left(\frac{(g24s + r\ (1-r)\ (g14s + g23s))}{2}\right) + (1-S) * 2 * x2S0 * x4S0;$$
$$g34S = S\left(\frac{(g34s + r\ (1-r)\ (g14s + g23s))}{2}\right) + (1-S) * 2 * x3S0 * x4S0;$$

Allele frequencies are not affected by recombination, but LD is.

Specifically, the recombination term is reduced by a factor (1-F) due to reduction effective selfing. Hence selfing can maintain LD, especially if negative due to stabilising selection.

### Initial population z0 = 0

Under this condition, ab is the best haplotypes and all other combinations are deleterious

Let’s start by looking at LD, as that also features in the analysis of allele frequencies:

We first assume that γA, γB and z0 are small compared to Sqrt[2Vs] and we rescale these parameters by Sqrt[2Vs]

In[ ]:=
```
δ° = Normal[Series[(g11S + (g12S + g13S + g14S)/2) (g44S + (g14S + g24S + g34S)/2) -
     (g22S + (g12S + g23S + g24S)/2) (g33S + (g13S + g23S + g34S)/2) /.
     Gtype2Htype /. PDtoX /. Ssub, {ξ, 0, 2}]] /. ξ → 1 // Simplify
```

Out[ ]=

$$-pA^2\,\gamma A\,\left(-pB\,(\gamma B+3\,F\,\gamma B)+pB^2\,(\gamma B+3\,F\,\gamma B)+(-1+F)\,(2+(-1+F)\,r)\,\gamma A\,\delta\right)+\frac{1}{2}\,\delta\,\Big(2-\gamma A^2-3\,F\,\gamma A^2-2\,\gamma A\,\gamma B-6\,F\,\gamma A\,\gamma B+8\,F\,pB\,\gamma A\,\gamma B-\gamma B^2-3\,F\,\gamma B^2+8\,F\,pB\,\gamma B^2+4\,pB^2\,\gamma B^2-4\,F\,pB^2\,\gamma B^2+2\,d0\,(1+F)\,(\gamma A+\gamma B-2\,pB\,\gamma B)+2\,\gamma A\,\gamma B\,\delta+6\,F\,\gamma A\,\gamma B\,\delta+(-1+F)\,r\,\big(2-\gamma A^2-\gamma B^2+2\,pB\,\gamma B^2+2\,F\,pB\,\gamma B^2+2\,pB^2\,\gamma B^2-2\,F\,pB^2\,\gamma B^2+2\,d0\,(\gamma A+\gamma B-2\,pB\,\gamma B)+2\,\gamma A\,\gamma B\,(-1+2\,(1+F)\,\delta)\big)\Big)+pA\,\gamma A\,\Big(pB^2\,(\gamma B+3\,F\,\gamma B)+\big(-2\,d0\,(1+F-r+F\,r)-r\,\gamma A+F^2\,r\,\gamma A+4\,F\,(\gamma A+\gamma B)\big)\,\delta+pB\,\gamma B\,(-1-4\,(-1+r)\,\delta+F\,(-3+4\,(-1+r)\,\delta))\Big)$$

In[ ]:=
```
Δδ[F_, r_, d0_, γA_, γB_, pA_, pB_, δ_] = Simplify[δ° - δ]
```

Out[ ]=

$$-\delta-pA^2\,\gamma A\,\left(-pB\,(\gamma B+3\,F\,\gamma B)+pB^2\,(\gamma B+3\,F\,\gamma B)+(-1+F)\,(2+(-1+F)\,r)\,\gamma A\,\delta\right)+\frac{1}{2}\,\delta\,\Big(2-\gamma A^2-3\,F\,\gamma A^2-2\,\gamma A\,\gamma B-6\,F\,\gamma A\,\gamma B+8\,F\,pB\,\gamma A\,\gamma B-\gamma B^2-3\,F\,\gamma B^2+8\,F\,pB\,\gamma B^2+4\,pB^2\,\gamma B^2-4\,F\,pB^2\,\gamma B^2+2\,d0\,(1+F)\,(\gamma A+\gamma B-2\,pB\,\gamma B)+2\,\gamma A\,\gamma B\,\delta+6\,F\,\gamma A\,\gamma B\,\delta+(-1+F)\,r\,\big(2-\gamma A^2-\gamma B^2+2\,pB\,\gamma B^2+2\,F\,pB\,\gamma B^2+2\,pB^2\,\gamma B^2-2\,F\,pB^2\,\gamma B^2+2\,d0\,(\gamma A+\gamma B-2\,pB\,\gamma B)+2\,\gamma A\,\gamma B\,(-1+2\,(1+F)\,\delta)\big)\Big)+pA\,\gamma A\,\Big(pB^2\,(\gamma B+3\,F\,\gamma B)+\big(-2\,d0\,(1+F-r+F\,r)-r\,\gamma A+F^2\,r\,\gamma A+4\,F\,(\gamma A+\gamma B)\big)\,\delta+pB\,\gamma B\,(-1-4\,(-1+r)\,\delta+F\,(-3+4\,(-1+r)\,\delta))\Big)$$

As the two derived alleles are deleterious when z0 = 0, we can assume that pA, pB << 1. pA and pB are thus of order O($\zeta$) and $\delta$ of $O(\zeta^2)$ as it is of the order of the product pA pB.

In[ ]:=
```
ΔδapproxOptimum[F_, r_, γA_, γB_, pA_, pB_, δ_] = FullSimplify[
  Normal[Series[Δδ[F, r, 0, γA, γB, pA ξ, pB ξ, δ ξ^2], {ξ, 0, 2}]] /. ξ → 1]
```

Out[ ]=

$$-((1+3\,F)\,pA\,pB\,\gamma A\,\gamma B)-\frac{1}{2}\,\left((1+3\,F)\,(\gamma A+\gamma B)^2+(-1+F)\,r\,\left(-2+(\gamma A+\gamma B)^2\right)\right)\,\delta$$

In[ ]:= $\text{Simplify}\left[-((1+3F)\,pA\,pB\,\gamma A\,\gamma B)-\frac{1}{2}\left((1+3F)\,(\gamma A+\gamma B)^2+(1-F)\,r\,\left(2-(\gamma A+\gamma B)^2\right)\right)\delta-\left\{-((1+3F)\,pA\,pB\,\gamma A\,\gamma B)-\frac{1}{2}\left((1+3F)\,(\gamma A+\gamma B)^2+(-1+F)\,r\,\left(-2+(\gamma A+\gamma B)^2\right)\right)\delta\right\}\right]$

Out[ ]= $\{0\}$

In[ ]:= $\text{Solve}[\Delta\delta\text{approxOptimum}[F, r, \gamma A, \gamma B, pA, pB, \delta] == 0, \delta]$

Out[ ]= $\left\{\left\{\delta\to\frac{2\,(1+3F)\,pA\,pB\,\gamma A\,\gamma B}{-\left((1+3F)\,(\gamma A+\gamma B)^2\right)-(-1+F)\,r\,\left(-2+(\gamma A+\gamma B)^2\right)}\right\}\right\}$

In[ ]:= $\text{Simplify}\left[-\frac{2\,(1+3F)\,pA\,pB\,\gamma A\,\gamma B}{(1+3F)\,(\gamma A+\gamma B)^2+(1-F)\,r\,\left(2-(\gamma A+\gamma B)^2\right)}-\left\{\frac{2\,(1+3F)\,pA\,pB\,\gamma A\,\gamma B}{-(1+3F)\,(\gamma A+\gamma B)^2-(-1+F)\,r\,\left(-2+(\gamma A+\gamma B)^2\right)}\right\}\right]$

Out[ ]= $\{0\}$

From this result , the first term shows that selection generates negative LD when the effect of the two alleles are aligned ($\gamma A\ \gamma B > 0$) and positive LD when two alleles have opposite effects ($\gamma A\ \gamma B < 0$) and that selfing reinforces this effect.

The second term shows that recombination breaks down LD, but less efficiently under selfing, term 1-F. Note that because we assume that $\gamma A$ and $\gamma B$ are small, $\left(2-(\gamma A+\gamma B)^2\right)$ is always positive.

This second term also shows that selection also reduces LD but at the order $\delta$, and this effect is amplified by selfing. The net effect of selfing on the second term depend on the recombination / selection balance.

In[ ]:= $\text{D}\left[(1+3F)\,(\gamma A+\gamma B)^2+(1-F)\,r\,\left(2-(\gamma A+\gamma B)^2\right), F\right]$

Out[ ]= $3\,(\gamma A+\gamma B)^2-r\,\left(2-(\gamma A+\gamma B)^2\right)$

In[ ]:= $\text{Solve}[\% == 0, r]\ //\ \text{Simplify}$

Out[ ]= $\left\{\left\{r\to-\frac{3\,(\gamma A+\gamma B)^2}{-2+\gamma A^2+2\,\gamma A\,\gamma B+\gamma B^2}\right\}\right\}$

Approximately, selfing slow downs the erosion of LD in the second term when recombination is stronger than selection: $r > 3\ (\gamma A+\gamma B)^2$.

Otherwise, selfing facilitates the erosion of LD. The first condition corresponds to QLE conditions and if so, $\delta$ at QLE reduces to:

In[ ]:= $\text{Normal}\left[\text{Series}\left[-\frac{2\,(1+3F)\,pA\,pB\,\gamma A\,\xi\,\gamma B\,\xi}{(1+3F)\,(\gamma A\,\xi+\gamma B\,\xi)^2+(1-F)\,r\,\left(2-(\gamma A\,\xi+\gamma B\,\xi)^2\right)}, \{\xi, 0, 2\}\right]\right]$

Out[ ]= $\frac{(1+3F)\,pA\,pB\,\gamma A\,\gamma B\,\xi^2}{(-1+F)\,r}$

$\delta\text{QLEapprox}[F\_, r\_, pA\_, pB\_, \gamma A\_, \gamma B\_] = -\frac{(1+3F)\,pA\,pB\,\gamma A\,\gamma B}{(1-F)\,r};$

We then applies the same approach to allelic frequencies

In[ ]:=
```
pA° = Simplify[
  Normal[Series[g12s/2 + g14s/2 + g22s + g23s/2 + g24s + g34s/2 + g44s /. Gtype2Htype /.
      PDtoX /. Ssub, {ξ, 0, 2}]] /. ξ → 1]
```

Out[ ]=

$$pA + \frac{1}{2}\left(-2\,(-1+F)\,pA^3\,\gamma A^2 + pA^2\,\gamma A\,((-1+5\,F)\,\gamma A + 4\,(1+F)\,pB\,\gamma B) + \gamma B\,(-2\,(1+3\,F)\,\gamma A + (-1-3\,F-2\,pB+2\,F\,pB)\,\gamma B)\,\delta - 2\,d0\,(1+F)\left(-pA\,\gamma A + pA^2\,\gamma A - \gamma B\,\delta\right) - pA\,\gamma A\,(\gamma A + 3\,F\,\gamma A + 4\,\gamma B\,(pB + F\,pB - 2\,F\,\delta))\right)$$

In[ ]:=
```
ΔpA[F_, d0_, γA_, γB_, pA_, pB_, δ_] = Simplify[pA° - pA]
```

Out[ ]=

$$\frac{1}{2}\left(-2\,(-1+F)\,pA^3\,\gamma A^2 + pA^2\,\gamma A\,((-1+5\,F)\,\gamma A + 4\,(1+F)\,pB\,\gamma B) + \gamma B\,(-2\,(1+3\,F)\,\gamma A + (-1-3\,F-2\,pB+2\,F\,pB)\,\gamma B)\,\delta - 2\,d0\,(1+F)\left(-pA\,\gamma A + pA^2\,\gamma A - \gamma B\,\delta\right) - pA\,\gamma A\,(\gamma A + 3\,F\,\gamma A + 4\,\gamma B\,(pB + F\,pB - 2\,F\,\delta))\right)$$

In[ ]:=
```
ΔpB[F_, d0_, γA_, γB_, pA_, pB_, δ_] = ΔpA[F, z0, γB, γA, pB, pA, δ]
```

Out[ ]=

$$\frac{1}{2}\left(-2\,(-1+F)\,pB^3\,\gamma B^2 + pB^2\,\gamma B\,(4\,(1+F)\,pA\,\gamma A + (-1+5\,F)\,\gamma B) + \gamma A\,((-1-3\,F-2\,pA+2\,F\,pA)\,\gamma A - 2\,(1+3\,F)\,\gamma B)\,\delta - 2\,(1+F)\,z0\left(-pB\,\gamma B + pB^2\,\gamma B - \gamma A\,\delta\right) - pB\,\gamma B\,(\gamma B + 3\,F\,\gamma B + 4\,\gamma A\,(pA + F\,pA - 2\,F\,\delta))\right)$$

Even after simplification the dynamics depends in a complex manner of the different parameters

In[ ]:=
```
ΔpAoptapprox[F_, γA_, γB_, pA_, pB_, δ_] =
 FullSimplify[Normal[Series[ΔpA[F, 0, γA, γB, pA ξ, pB ξ, δ ξ^2], {ξ, 0, 2}]] /. ξ → 1]
```

Out[ ]=

$$\frac{1}{2}\,(pA\,\gamma A\,((-1-pA+F\,(-3+5\,pA))\,\gamma A - 4\,(1+F)\,pB\,\gamma B) - (1+3\,F)\,\gamma B\,(2\,\gamma A + \gamma B)\,\delta)$$

Relatively to a panmictic population selfing has an effect independent of r and γ on opposed alleles whereas the effect on LD on aligned alleles depends on the balance between recombination and selection.
When recombination is high compared to selection, the effect on opposed and aligned alleles are very similar
Intermediate selfing reduces LD (F<2/3 of F < 2/3 * f(γ,r)) but high selfing amplifies it.
Compared to the results above for any pA and pB, the diffrence here comes from the fact that selfing reduces the frequency of deleterious alleles (purging) which reduces the absolute LD value.
LD increases again when the effect of reducing recombination overwhelms the purging effect.

### Shift to a new optimum z0 ≠ 0

In[ ]:= 
```
z0 / Sqrt[Vs]
```
Out[ ]=

$$\frac{z0}{\sqrt{Vs}}$$

Without loss of generality we will assume that z0 > 0. We still work with the scaled parameter so with $d0 = \frac{z0}{\sqrt{Vs}}$

If we assume that |γA|, |γB| << d0, we can expand the expression as follows:

In[ ]:= 
```
FullSimplify[Normal[Series[ΔpA[F, d0, γA ξ, γB ξ, pA , pB , δ], {ξ, 0, 1}]] /. ξ → 1]
```
Out[ ]=

$$d0\ (1 + F)\ (-((-1 + pA)\ pA\ \gamma A) + \gamma B\ \delta)$$

In[ ]:= 
```
Collect[ -d0 (1 + F) ((-1 + pA) pA γA - γB δ), δ]
```
Out[ ]=

$$-d0\ (1 + F)\ (-1 + pA)\ pA\ \gamma A + d0\ (1 + F)\ \gamma B\ \delta$$

The first term corresponds to direct selection and depends on the sign of γA. As expected when γA > 0 alleles are positively selected and when γA < 0 they are negatively selected.
Selfing increases selection by (1+F). But if we consider the effect of drift, Ne = N/(1+F), the two effects cancels out. So if we neglect associations the initial dynamics should be unaffected by selfing.
The second term correspond to linked selection and is always of the opposite sign of γA because γA γB δ is always negative. So initial associations slow down the response to selection.
As selfing magnifies LD, selfing should reduce the response to selection because of genetic associations.

We can then express the recursion on LD

In[ ]:= 
```
Δδshift[F_, r_, d0_, γA_, γB_, pA_, pB_, δ_] =
  FullSimplify[Normal[Series[Δδ[F, r, d0, γA ξ, γB ξ, pA, pB, δ], {ξ, 0, 1}]] /. ξ → 1]
```
Out[ ]=

$$(-r + F\ r - 2\ d0\ pA\ (1 + F + (-1 + F)\ r)\ \gamma A + d0\ (1 + F)\ (\gamma A + \gamma B - 2\ pB\ \gamma B) + d0\ (-1 + F)\ r\ (\gamma A + \gamma B - 2\ pB\ \gamma B))\ \delta$$

Simplification by hand. We note re = r (1 - F) and κ = (1-2 pA) γA+(1-2 pB) γB

In[ ]:= 
```
FullSimplify[
   Δδshift[F, r, d0, γA, γB, pA, pB, 1] - Δδshift[F, 0, d0, γA, γB, pA, pB, 1]] +
  FullSimplify[Δδshift[F, 0, d0, γA, γB, pA, pB, 1]]
```
Out[ ]=

$$d0\ (1 + F)\ (\gamma A - 2\ pA\ \gamma A + \gamma B - 2\ pB\ \gamma B) - (-1 + F)\ r\ (-1 + d0\ (-1 + 2\ pA)\ \gamma A + d0\ (-1 + 2\ pB)\ \gamma B)$$

We also assume weak selection so d0 (-1 + 2 pA) γA + d0 (-1 + 2 pB) γB << 1, which leads to the equation in the main text

In[ ]:= 
```
Δδshift[F_, r_, κ_] = δ Simplify[d0 (1 + F) (κ) - (-1 + F) r (-1)]
```
Out[ ]=

$$\delta\ ((-1 + F)\ r + d0\ (1 + F)\ \kappa)$$
