## Supplementary File S2 for "Polygenic selection to a changing optimum under self–fertilisation": S2_File.pdf

#### S2 File: Dynamics at a locus underlying a quantitative trait under partial selfing

Matthew Hartfield and Sylvain Glémin

*In this notebook we analyse the dynamics of a single (focal) locus underlying a quantitative trait under partial selfing.*

#### General model

We consider a quantitative trait affected by $n$ bi-allelic loci with additive effect:
$A_i\, A_i$: $2\,\alpha_i$
$A_i\, a_i$: $\alpha_i$
$a_i\, a_i$: 0
We note $x_i$ the frequency of allele $A_i$

The mean phenotypic effect of locus $i$ is $2\,\alpha_i\, x_i$

In[∘ ]:=
```
Simplify[2 αi (xi^2 + F xi (1 - xi)) + αi 2 xi (1 - xi) (1 - F)]
```

Out[∘ ]=

$2\, x_i\, \alpha_i$

We also note $z$ the mean phenotype of the population with $z = \Sigma_i\, 2\,\alpha_i\, x_i$

In what follows we consider a single focal locus and the $n$ - 1 other loci are considered as the background.

The mean phenotypic effect of the background is thus $z_i = z - 2\,\alpha_i\, x_i$
If we neglect associations, the background is the same for the two alleles, hence the phenotype of the three genotypes are:
$A_i\, A_i$: $2\,\alpha_i + z - 2\,\alpha_i\, x_i$
$A_i\, a_i$: $\alpha_i + z - 2\,\alpha_i\, x_i$
$a_i\, a_i$: $z - 2\,\alpha_i\, x_i$
In the more general case we can note that allele $A_i$ is associated with the background $z_i + \beta_{A,i}$ and $a_i$ with the background $z_i + \beta_{a,i}$

As we have $\beta_{A,i}\, x_i + \beta_{a,i}(1 - x_i) = 0$.

We can thus write:
$\beta_{A,i} = \beta_i(1 - x_i)$
$\beta_{a,i} = -\beta_i\, x_i$

So the mean phenotypic effect of the three genotypes can be written as:

$A_i\,A_i$: $2\,\alpha_i - 2\,\alpha_i\,x_i + 2\,\beta_i(1 - x_i)$

$A_i\,a_i$: $\alpha_i - 2\,\alpha_i\,x_i + \beta_i(1 - x_i) - \beta_i\,x_i$

$a_i\,a_i$: $-2\,\alpha_i\,x_i - 2\,\beta_i\,x_i$

Then we can apply the classical gaussian selection model.

We note Vs = $\omega^2$ + Ve, with $\omega$ is the standard deviation of the gaussian fitness function and Ve the environmental variance.

In what follows we remove the subscript *i* for simplicity.

```
In[ ]:= zAA = 2 α + (z - 2 α x) + 2 β (1 - x);
        zAa = α + (z - 2 α x) + β (1 - x) - β x;
        zaa = 0 + (z - 2 α x) - 2 β x;
        zm = Simplify[(x^2 + F x (1 - x)) zAA + 2 x (1 - x) (1 - F) zAa + ((1 - x)^2 + F x (1 - x)) zaa];
        w[z_] = E^(-(z - z0)^2/(2 Vs));
        Wm = Simplify[
           (x^2 + F x (1 - x)) w[zAA] + 2 x (1 - x) (1 - F) w[zAa] + ((1 - x)^2 + F x (1 - x)) w[zaa]];
```

The change in allele frequency due to selection is given by:

```
In[ ]:= dx = Simplify[((x^2 + F x (1 - x)) w[zAA] + x (1 - x) (1 - F) w[zAa])/Wm - x]
```

Out[ ]=

$$-x + \left(e^{-\frac{(-z+z0+(-1+2x)(\alpha+\beta))^2}{2\,Vs}}(-1+F)(-1+x)\,x + e^{-\frac{(-z+z0+2(-1+x)(\alpha+\beta))^2}{2\,Vs}}\,x\,(F+x-F\,x)\right) \Big/ \left(2\,e^{-\frac{(-z+z0+(-1+2x)(\alpha+\beta))^2}{2\,Vs}}(-1+F)(-1+x)\,x - e^{-\frac{(-z+z0+2x(\alpha+\beta))^2}{2\,Vs}}(-1+x)(1+(-1+F)\,x) + e^{-\frac{(-z+z0+2(-1+x)(\alpha+\beta))^2}{2\,Vs}}\,x\,(F+x-F\,x)\right)$$

We note $\mathcal{D}$ = z0 - z, the distance of the mean trait to the optimum

We can also express z, z0, $\beta$ and $\alpha$ in $\sqrt{Vs}$ unit to simplify the expression.

The re-scaled parameters are written with °

```
In[ ]:= Δx[F_, D°_, α°_, β°_, x_] = Simplify[
          dx /. z → z0 - D /. {D → D° Sqrt[Vs], β → β° Sqrt[Vs], α → α° Sqrt[Vs]}, Vs > 0]
```

Out[ ]=

$$-x + \left(e^{-\frac{1}{2}(\mathcal{D}^\circ+(-1+2x)(\alpha^\circ+\beta^\circ))^2}(-1+F)(-1+x)\,x + e^{-\frac{1}{2}(\mathcal{D}^\circ+2(-1+x)(\alpha^\circ+\beta^\circ))^2}\,x\,(F+x-F\,x)\right) \Big/ \left(2\,e^{-\frac{1}{2}(\mathcal{D}^\circ+(-1+2x)(\alpha^\circ+\beta^\circ))^2}(-1+F)(-1+x)\,x - e^{-\frac{1}{2}(\mathcal{D}^\circ+2x(\alpha^\circ+\beta^\circ))^2}(-1+x)(1+(-1+F)\,x) + e^{-\frac{1}{2}(\mathcal{D}^\circ+2(-1+x)(\alpha^\circ+\beta^\circ))^2}\,x\,(F+x-F\,x)\right)$$

### Equilibrium ($\mathcal{D}$ = 0)

#### Allele frequency

We assume weak selection, so $\alpha°$, $\beta°$ << 1. We thus obtain a simple expression for the change in allele frequency:

In[ ]:=
```
Δxsel0[F_, α°_, β°_, x_] =
  Simplify[Normal[Series[Δx[F, 0 , α° ξ, β° ξ, x], {ξ, 0, 2}]] /. ξ → 1]
```
Out[ ]=

$$-\frac{1}{2}\,(1+3\,F)\,x\,\left(1-3\,x+2\,x^2\right)\,(\alpha^\circ+\beta^\circ)^2$$

We also consider symmetrical mutation

In[ ]:=
```
Δmut[u_, x_] = Simplify[ -u x + u (1 - x)];
```

So we can get the frequency at equilibrium:

In[ ]:=
```
xeq[F_, α°_, β°_, u_] = Simplify[x /. Solve[Δxsel0[F, α°, β°, x] + Δmut[u, x] == 0, x]]
```
Out[ ]=

$$\left\{\frac{1}{2},\ \frac{\alpha^\circ+3\,F\,\alpha^\circ+\beta^\circ+3\,F\,\beta^\circ-\sqrt{1+3\,F}\,\sqrt{-8\,u+(1+3\,F)\,(\alpha^\circ+\beta^\circ)^2}}{2\,(1+3\,F)\,(\alpha^\circ+\beta^\circ)},\right.$$
$$\left.\frac{\alpha^\circ+3\,F\,\alpha^\circ+\beta^\circ+3\,F\,\beta^\circ+\sqrt{1+3\,F}\,\sqrt{-8\,u+(1+3\,F)\,(\alpha^\circ+\beta^\circ)^2}}{2\,(1+3\,F)\,(\alpha^\circ+\beta^\circ)}\right\}$$

And for low mutation rate

In[ ]:=
```
Simplify[Series[xeq[F, α°, β°, u], {u, 0, 1}], {α° + β° > 0}]
Simplify[Series[xeq[F, α°, β°, u], {u, 0, 1}], {α° + β° < 0}]
```
Out[ ]=

$$\left\{\frac{1}{2},\ \frac{2\,u}{(1+3\,F)\,(\alpha^\circ+\beta^\circ)^2}+O[u]^2,\ 1-\frac{2\,u}{(1+3\,F)\,(\alpha^\circ+\beta^\circ)^2}+O[u]^2\right\}$$

Out[ ]=

$$\left\{\frac{1}{2},\ 1-\frac{2\,u}{(1+3\,F)\,(\alpha^\circ+\beta^\circ)^2}+O[u]^2,\ \frac{2\,u}{(1+3\,F)\,(\alpha^\circ+\beta^\circ)^2}+O[u]^2\right\}$$

Which we can write with the initial, unscaled, parameters.

In[ ]:=
```
FullSimplify[{2 u/((1 + 3 F) (α° + β°)^2), 1 - 2 u/((1 + 3 F) (α° + β°)^2)} /.
   {α° → α / Sqrt[Vs], β° → β / Sqrt[Vs]}, Vs > 0]
```
Out[ ]=

$$\left\{\frac{2\,u\,Vs}{(1+3\,F)\,(\alpha+\beta)^2},\ 1-\frac{2\,u\,Vs}{(1+3\,F)\,(\alpha+\beta)^2}\right\}$$

There are two symmetrical equilibria, close to 0 or 1, and one central equilibrium at 1/2. The stability of equilibria depends on the sign of the derivative of Δx:

In[ ]:=
```
FullSimplify[D[Δxsel0[F, α°, β°, x] + Δmut[u, x], x] /. x → xeq[F, α°, β°, u]]
```
Out[ ]=

$$\left\{-2\,u+\frac{1}{4}\,(1+3\,F)\,(\alpha^\circ+\beta^\circ)^2,\ 4\,u-\frac{1}{2}\,(1+3\,F)\,(\alpha^\circ+\beta^\circ)^2,\ 4\,u-\frac{1}{2}\,(1+3\,F)\,(\alpha^\circ+\beta^\circ)^2\right\}$$

The 1/2 equilibrium is stable only if selection is weak compare to mutation, that is:

$$(\beta^\circ+\alpha^\circ)^2<\frac{8\,u}{1+3\,F}$$

Or with initial parameters

$$(\beta+\alpha)^2<\frac{8\,u\,Vs}{1+3\,F}$$

When $\beta = 0$ we retrieve the result of Abu-Awad and Roze (2018).
As stabilizing selection generates negative LD (see also the two-locus model), $\alpha$ and $\beta$ have opposite sign, so genetic associations leads to higher equilibrium frequencies than predicted by single locus theory.

#### Variance components

##### Derivations

We can also obtain the different variance component at equilibrium.
According to Bulmer (1976), the total genetic variance can be decomposed into:
- The genic variance, $V_g$, corresponding to Hardy–Weinberg expectations and no linkage
- The inbreeding variance, $V_I$, corresponding to the departure from Hardy-Weinberg expactations at each locus
- The linkage covariance, $C_{LD}$, which takes interactions among loci into account

The genic and inbreeding variance are readily obtained from the sum of each locus contribution:

```
In[ ]:= Vg[F_, α_, β_, x_] = Simplify[x^2 (2 α)^2 + 2 x (1 - x) (α)^2 + (1 - x)^2 (0)^2]
        VI[F_, α_, β_, x_] =
         Simplify[(F x (1 - x)) (2 α)^2 + -2 F x (1 - x) (α)^2 + (F x (1 - x)) (0)^2]
```

Out[ ]=

$$2\,x\,(1+x)\,\alpha^2$$

Out[ ]=

$$-2\,F\,(-1+x)\,x\,\alpha^2$$

Which can be evaluated at equilibrium and assuming mutation rate is small.
For low frequency equilibrium:

```
In[ ]:= Simplify[Normal[Series[Vg[F, α, β, (2 u Vs)/((1 + 3 F) (α + β)^2)], {u, 0, 1}]]]
        Simplify[Normal[Series[VI[F, α, β, (2 u Vs)/((1 + 3 F) (α + β)^2)], {u, 0, 1}]]]
```

Out[ ]=

$$\frac{4\,u\,Vs\,\alpha^2}{(1+3\,F)\,(\alpha+\beta)^2}$$

Out[ ]=

$$\frac{4\,F\,u\,Vs\,\alpha^2}{(1+3\,F)\,(\alpha+\beta)^2}$$

For intermediate frequency equilibrium (1/2):

In[ ]:=
```
Simplify[Normal[Series[Vg[F, α, β, 1 / 2], {u, 0, 1}]]]
Simplify[Normal[Series[VI[F, α, β, 1 / 2], {u, 0, 1}]]]
```
Out[ ]=

$$\frac{3\,\alpha^2}{2}$$

Out[ ]=

$$\frac{1}{2}\,F\,(\alpha+\beta)^2$$

If there is no locus at intermediate equilibrium:

$V_g = \sum_{i=1}^{n} \frac{4\,u_i\,V_s}{1+3\,F}\,\frac{\alpha_i^2}{(\alpha_i+\beta_i)^2}$

$V_I = \sum_{i=1}^{n} \frac{4\,F\,u_i\,V_s}{1+3\,F}\,\frac{\alpha_i^2}{(\alpha_i+\beta_i)^2}$

Further assuming that mutation rates and phenotypic effects are independent (or constant mutation rate) then:

$$V_g = \frac{4\,U\,V_s}{1+3\,F}\,\frac{1}{n}\sum_{i=1}^{n}\frac{\alpha_i^2}{(\alpha_i+\beta_i)^2}$$

$$V_I = \frac{4\,F\,U\,V_s}{1+3\,F}\,\frac{1}{n}\sum_{i=1}^{n}\frac{\alpha_i^2}{(\alpha_i+\beta_i)^2}$$

The covariance due to LD can also be computed as the covariance of phenotypic effect of the locus and the background.

Adapting Bulmer (1976), we have:

$C_{\text{LD}} = \sum_{i=1}^{n} \sum_{j\neq i} \text{Cov}(i,\,j)\,\alpha_i\,\alpha_j$

where Cov($i$, $j$) is the covariance between the number of allele $A$ at locus $i$ and $j$. Note that Bulmer considered that alleles contribute either 0 or 1 to the trait.

Note that $\text{Cov}(i,\,j)\,\alpha_i\,\alpha_j$ can be written as:

$$2\,\alpha_i\,f(A_i\,A_i)\left(f(A_j\,A_j \mid A_i\,A_i)\,2\,\alpha_j + f(A_j\,a_j \mid A_i\,A_i)\,\alpha_j + f(a_j\,a_j \mid A_i\,A_i)*0\right) +$$
$$\alpha_i\,f(A_i\,a_i)\left(f(A_j\,A_j \mid A_i\,a_i)\,2\,\alpha_j + f(A_j\,a_j \mid A_i\,A_i)\,\alpha_j + f(a_j\,a_j \mid A_i\,A_i)*0\right) +$$
$$0*f(a_i\,a_i)\left(f(A_j\,A_j \mid a_i\,a_i)\,2\,\alpha_j + f(A_j\,a_j \mid a_i\,a_i)\,\alpha_j + f(a_j\,a_j \mid a_i\,a_i)*0\right)$$

where $f(G_j \mid G_i)$ means the frequency of genotype $G_j$ knowing genotype $G_i$. Abu Awad and Roze expressed these quantities with a set of genetic associations, for which they seek for a set of recursion equations.

With our formalism, they are encapsulated in the $\beta$ parameters such that:

$2\,\beta_{A,i} = 2\,\beta_i(1-x_i) = \sum_{j\neq i}\left(f(A_j\,A_j \mid A_i\,A_i)\,2\,\alpha_j + f(A_j\,a_j \mid A_i\,A_i)\,\alpha_j + f(a_j\,a_j \mid A_i\,A_i)*0\right)$

$\beta_{A,i} + \beta_{a,i} = \beta_i(1-2\,x_i) = \sum_{j\neq i}\left(f(A_j\,A_j \mid A_i\,a_i)\,2\,\alpha_j + f(A_j\,a_j \mid A_i\,a_i)\,\alpha_j + f(a_j\,a_j \mid A_i\,a_i)*0\right)$

$2\,\beta_{a,i} = -2\,\beta_i\,x_i = \sum_{j\neq i}\left(f(A_j\,A_j \mid a_i\,a_i)\,2\,\alpha_j + f(A_j\,a_j \mid a_i\,a_i)\,\alpha_j + f(a_j\,a_j \mid a_i\,a_i)*0\right)$

So we can write:

$C_{\text{LD}} = \sum_{i=1}^{n} f(A_i\,A_i)\,4\,\alpha_i\,\beta_i(1-x_i) + f(A_i\,a_i)\,2\,\alpha_i\,\beta_i(1-2\,x_i) + f(a_i\,a_i)*0*(-2\,\beta_i\,x_i)$

For one locus:

In[ ]:=
```
Simplify[Normal[Series[Cld[F, α, β, (2 u Vs)/((1 + 3 F) (α + β)^2)], {u, 0, 1}]]]
```

Out[ ]=

$$\frac{4\ (1+F)\ u\ Vs\ \alpha\ \beta}{(1+3\ F)\ (\alpha+\beta)^2}$$

Summing over the *n* loci:

$$C_{LD} = \frac{4\,(1+F)\,U\,V_s}{1+3\,F}\,\frac{1}{n}\sum_{i=1}^{n}\frac{\alpha_i\,\beta_i}{(\alpha_i+\beta_i)^2}$$

Finally, by summing the three components we obtain the total genetic variance.
For one locus:

In[ ]:=
```
VG[F_, Vs_, α_, β_, u_] =
  FullSimplify[(4 u Vs α^2)/((1 + 3 F) (α + β)^2) + (4 F u Vs α^2)/((1 + 3 F) (α + β)^2) + (4 (1 + F) u Vs β α)/((1 + 3 F) (α + β)^2)]
```

Out[ ]=

$$\frac{4\ (1+F)\ u\ Vs\ \alpha}{(1+3\ F)\ (\alpha+\beta)}$$

And summed over all loci:

$$V_G = \frac{4\,(1+F)\,U\,V_s}{1+3\,F}\sum_{i=1}^{n}\frac{\alpha_i}{\alpha_i+\beta_i}$$

Note that because the total variance must be positive, $\frac{\alpha_i}{\alpha_i+\beta_i} > 0$. As $\alpha_i$ and $\beta_i$ are opposite sign it implies that $|\beta_i| < |\alpha_i|$

#### Interpretation

The expressions for variance components are not closed but they provide useful insight on the effect of genetic associations, encapsullated in $\beta_i$s, on variance decomposition.

- When $\beta_i = 0$ we retrieve the house of card expression with selfing as in Abu-Awad and Roze 2018 with $C_{LD} = 0$
- As $\alpha_i$ and $\beta_i$ are of opposite sign, associations increase the genic and inbreeding variance (because selection is reduced) and make $C_{LD}$ more negative. The overall effect is to increase genic variance compared to no associations
- Selfing has two opposite effects: increasing purging, which reduces the genetic variance but increasing associations, which increased the genetic variance. However, we can't determine the net effect, which would require to express $\sum_{i=1}^{n}\frac{\alpha_i}{\alpha_i+\beta_i}$ as a function of *F*.

### Shift to a new optimum

In this part we assume that the population is away from equilibrium. Without loss of generality we assume that d0 >0.

#### Change in allele frequency

As above we assume weak selection, so $\gamma°, \beta° << 1$. We also assume that $\mathcal{D}°<<1$, which means that the shift to the new optimum didn't generate a strong drop in fitness.

In[ ]:=
```
Δxsel[F_, 𝒟°_, α°_, β°_, x_] =
 FullSimplify[Normal[Series[Δx[F, 𝒟°, α° ξ, β° ξ, x], {ξ, 0, 2}]] /. ξ → 1]
```

Out[ ]=

$$\frac{1}{2}(-1+x)\,x\,(\alpha°+\beta°)\left(-2(1+F)\mathcal{D}°-(1+3F)(-1+2x)(\alpha°+\beta°)+(1+3F)(-1+2x)\mathcal{D}°^2(\alpha°+\beta°)\right)$$

It can be re-written as the sum of two terms. The first term corresponds to pure directional selection and the second to stabilising selection

In[ ]:=
```
Simplify[1/2 (-1 + x) x (α° + β°) (-2 𝒟° (1 + F))] + Simplify[
  1/2 (-1 + x) x (β° + α°) (-(1 + 3 F) (-1 + 2 x) (β° + α°) + 𝒟°^2 (1 + 3 F) (-1 + 2 x) (β° + α°))]
```

Out[ ]=

$$-(1+F)(-1+x)\,x\,\mathcal{D}°(\alpha°+\beta°)+\frac{1}{2}(1+3F)(-1+x)\,x\,(-1+2x)\left(-1+\mathcal{D}°^2\right)(\alpha°+\beta°)^2$$

This can be rewritten with the unscaled parameters

In[ ]:=
```
Simplify[1/2 (-1 + x) x (α° + β°) (-2 𝒟° (1 + F)) /.
   {α° → α/Sqrt[Vs], β° → β/Sqrt[Vs], 𝒟° → 𝒟/Sqrt[Vs]}, Vs > 0] + Simplify[1/2 (-1 + x)
   x (β° + α°) (-(1 + 3 F) (-1 + 2 x) (β° + α°) + 𝒟°^2 (1 + 3 F) (-1 + 2 x) (β° + α°)) /.
   {α° → α/Sqrt[Vs], β° → β/Sqrt[Vs], 𝒟° → 𝒟/Sqrt[Vs]}, Vs > 0]
```

Out[ ]=

$$-\frac{(1+F)(-1+x)\,x\,\mathcal{D}\,(\alpha+\beta)}{Vs}-\frac{(1+3F)(-1+x)\,x\,(-1+2x)\left(Vs-\mathcal{D}^2\right)(\alpha+\beta)^2}{2\,Vs^2}$$

If we neglect LD we obtain an expression equivalent to the one of Hayward and Sella 2022 with the additional effect of selfing

In[ ]:=
```
% /. β → 0
```

Out[ ]=

$$-\frac{(1+F)(-1+x)\,x\,\mathcal{D}\,\alpha}{Vs}-\frac{(1+3F)(-1+x)\,x\,(-1+2x)\left(Vs-\mathcal{D}^2\right)\alpha^2}{2\,Vs^2}$$

#### Change in the phenotype

The phenotypic change can be expressed in term of the distance to the optimum. Following Hayward and Sella 2022, we can write:

$$E[\Delta\mathcal{D}] = -\sum_{i=1}^{n} E[2\,\alpha_i\,\Delta x_i]$$

$$= -\frac{\mathcal{D}}{Vs}\sum_{i=1}^{n}\left(2\,\alpha_i^2(1+F)\,x_i(1-x_i)\;+2\,\alpha_i\,\beta_i(1+F)\,(1-x_i)\,x_i\right)+$$

$$\frac{(Vs-\mathcal{D}^2)}{Vs^2}\,(1+3F)\sum_{i=1}^{n}x_i\,(1-x_i)\left(\frac{1}{2}-x_i\right)\alpha_i\,(\alpha_i+\beta_i)^2$$

The first term we recognise the expressions for the variance components. In the second term, we need to introduce notations for third order moments

$\mu_3 = \sum_{i=1}^{n} x_i (1 - x_i) \left(\frac{1}{2} - x_i\right) \alpha_i^3$
$\nu_{2,1} = \sum_{i=1}^{n} x_i (1 - x_i) \left(\frac{1}{2} - x_i\right) \alpha_i^2 \beta_i$
$\nu_{1,2} = \sum_{i=1}^{n} x_i (1 - x_i) \left(\frac{1}{2} - x_i\right) \alpha_i \beta_i^2$

This leads to the expression in the main text

$$E[\Delta \mathcal{D}] = -\sum_{i=1}^{n} E[2\, \alpha_i\, \Delta x_i]$$

$$= -\frac{\mathcal{D}}{\text{Vs}} \sum_{i=1}^{n} \left((1 + F)\, V_g + C_{\text{LD}}\right) + \frac{\left(\text{Vs} - \mathcal{D}^2\right)}{\text{Vs}^2} (1 + 3\, F) (\mu_3 + 2\, \nu_{2,1} + \nu_{1,2})$$
