## Supplementary File S3 for "Polygenic selection to a changing optimum under self–fertilisation"

### Supplementary Figures

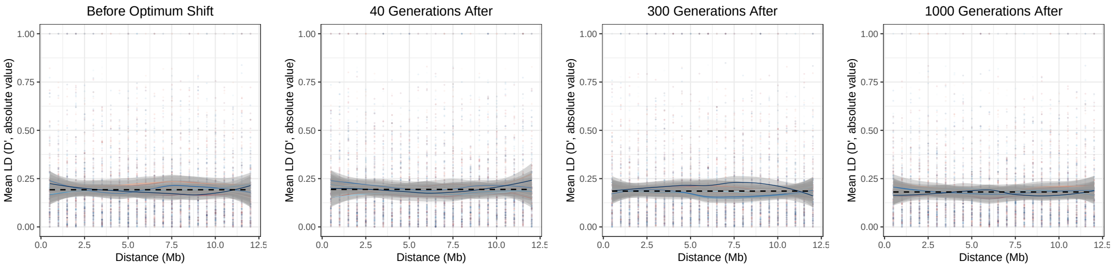


(a)

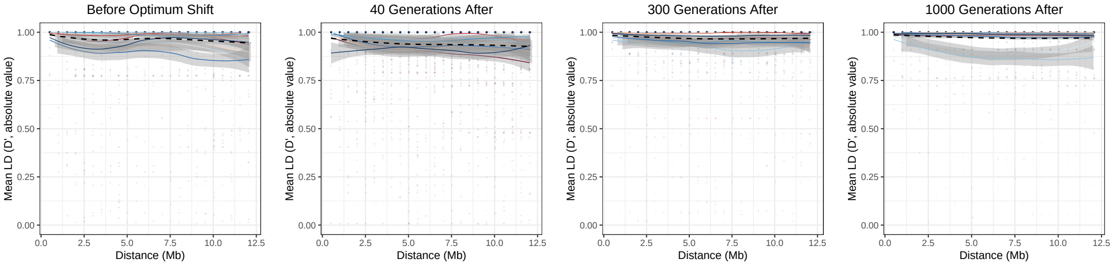


(b)

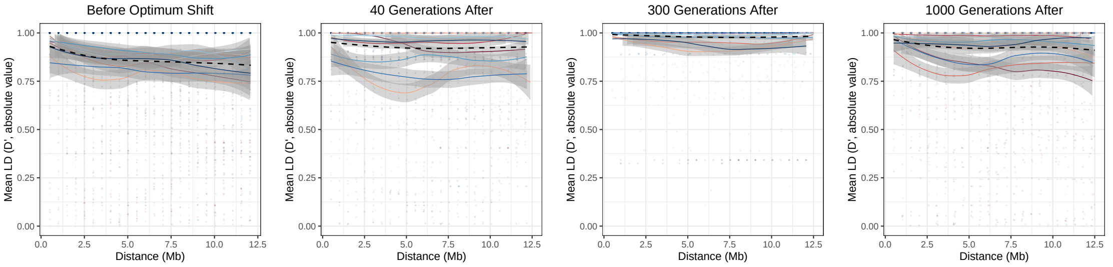


(c)

Figure A: As Figure 5 in the main text but plotting $|D'|$ as a function of distance.

### Gradual optimum shift, no background selection

In natural populations, it is also possible for organisms to experience a gradual optimum shift. This can occur if, for example, the environment changes slowly due to changes in the local environment or climate. This model also reflects cases where an organism is slowly expanding its range into new regions. A gradual optimum shift could also affect the LD structure in highly selfing species, due to the continual need to adapt over time. We hence ran simulations with a gradually–changing optimum to determine how this affected the nature of polygenic adaptation in selfing species, focusing only on the non–pleiotropic case.

Adaptation takes longer in line with the gradual change in optimum value (Figure B(a)). There is a noticeable dip in mean fitness during this adaptation phase; we also observe a rise in the variance in fitness. After 100 generations, we observe similar results to the instant–optimum–shift case, with high self–fertilising exhibiting the highest mean fitness and lowest inbreeding depression and fitness variance. Variance measurements (Figure B(b)) and haplotype plots (Figure B(b)) are qualitatively similar to the instant–shift case. Mean LD values appear constant over time as in the instant–shift case, but there's an appearance in high–LD cases later on at 1,000 generations (Figure D).

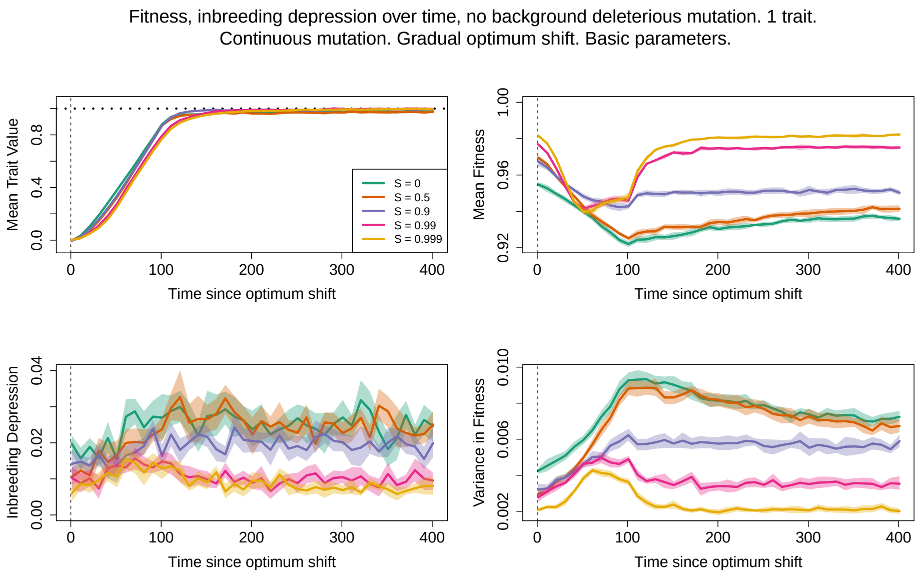


(a)

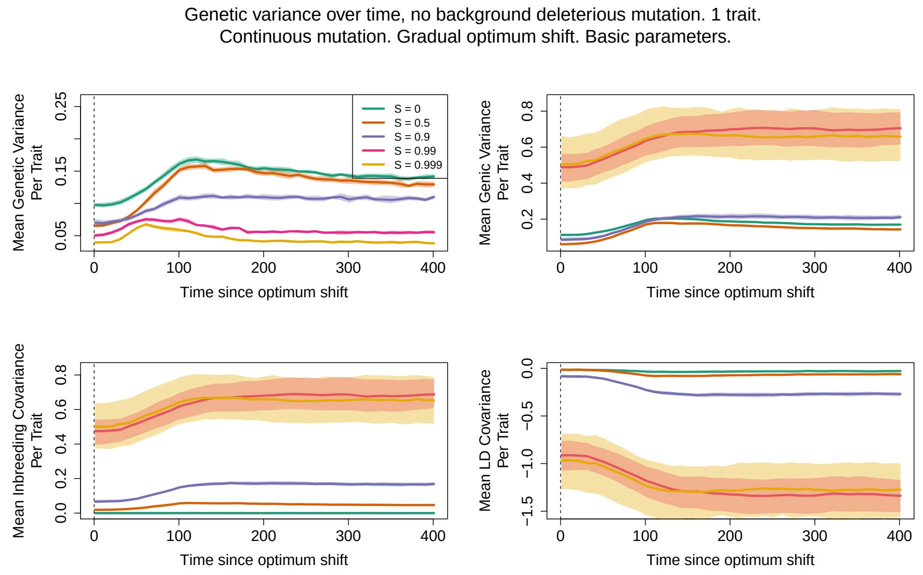


(b)

Figure B: (a) Mean fitness, (b) variance measurements where there is a gradual change in the fitness optimum and pleiotropy is absent.

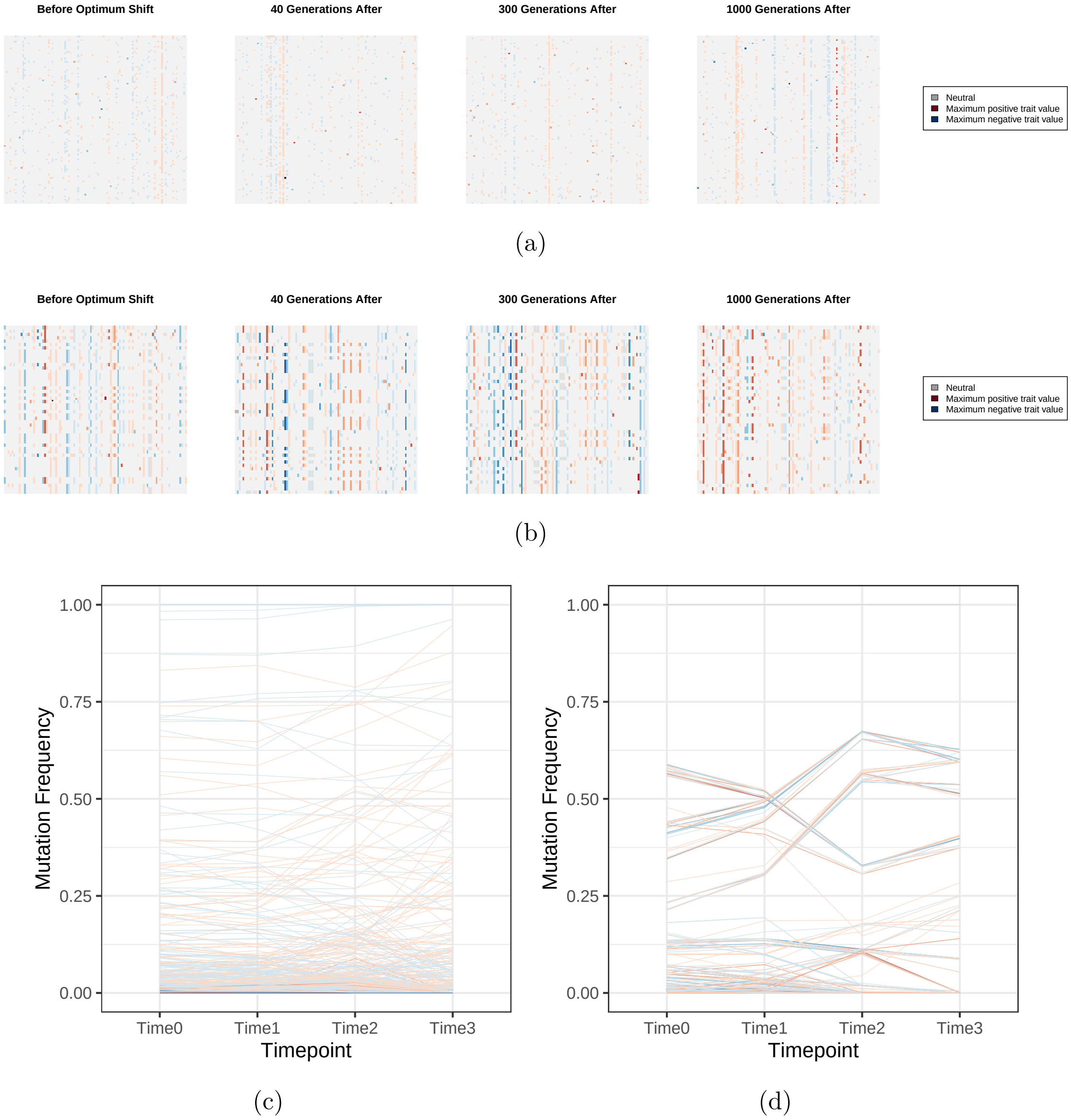


Figure C: (a), (b) Haplotype plots and (c), (d) following a gradual change in optimum shift when mutations are not pleiotropic. Populations are either (a) outcrossing or (b) 99.9% self–fertilising.

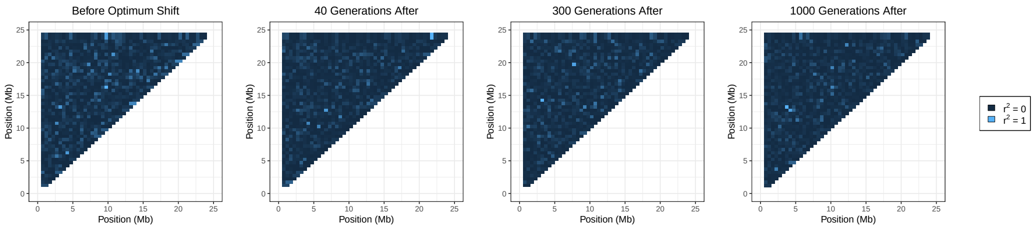


(a)

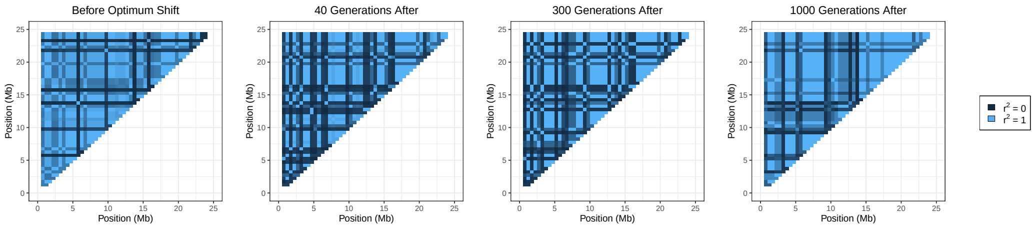


(b)

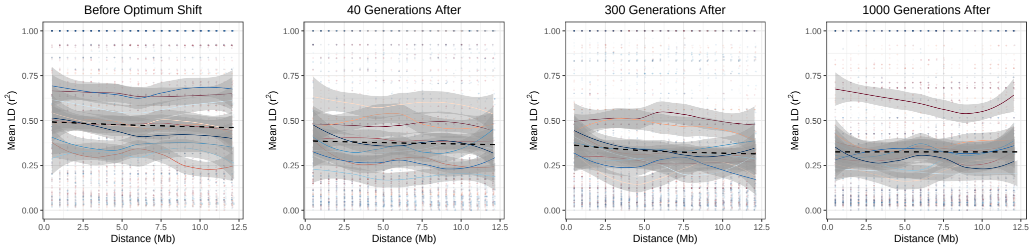


(c)

Figure D: (a), (b) LD heatmaps and (c) LD decay following a gradual change in optimum shift when mutations are not pleiotropic. Populations are either (a) outcrossing or (b), (c) 99.9% self–fertilising.

### Results with a larger population size

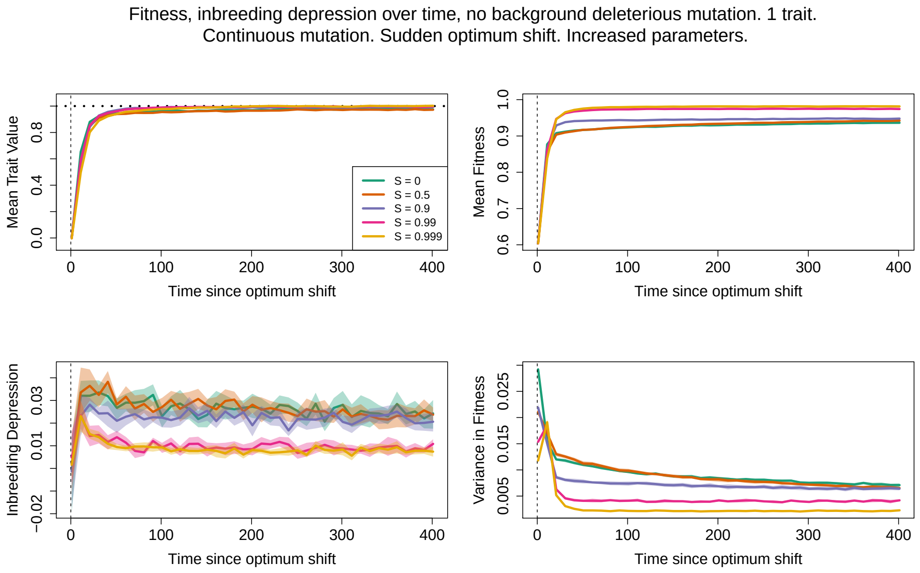


(a)

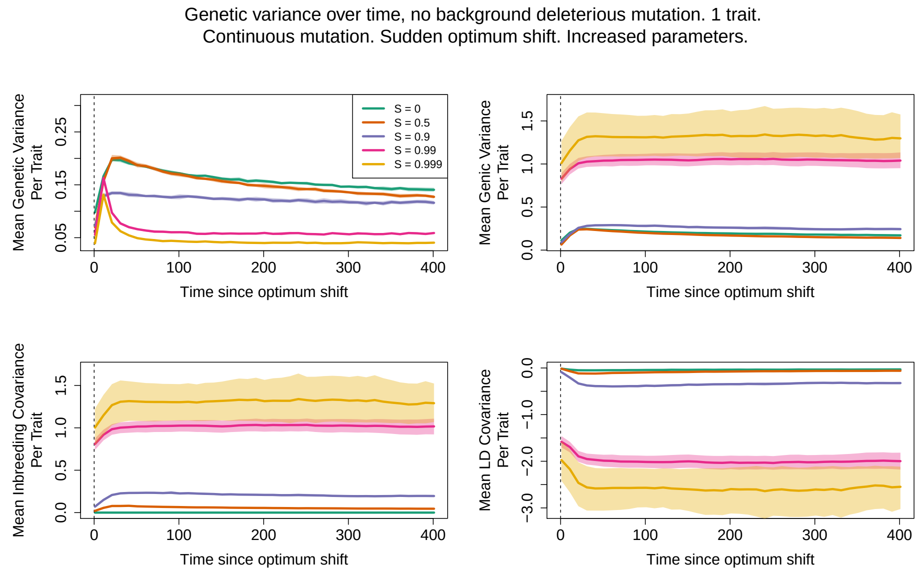


(b)

Figure E: (a) Mean fitness, (b) variance measurements for a population size $N = 10,000$.

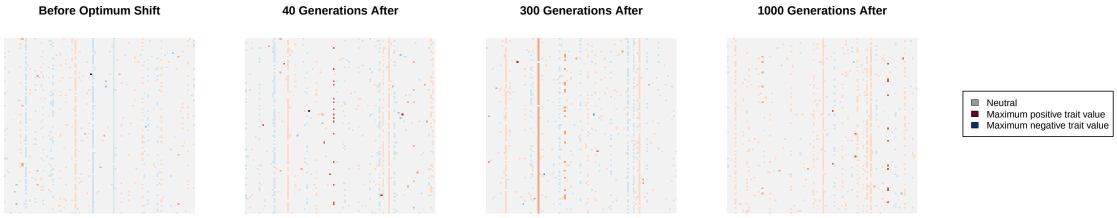


(a)

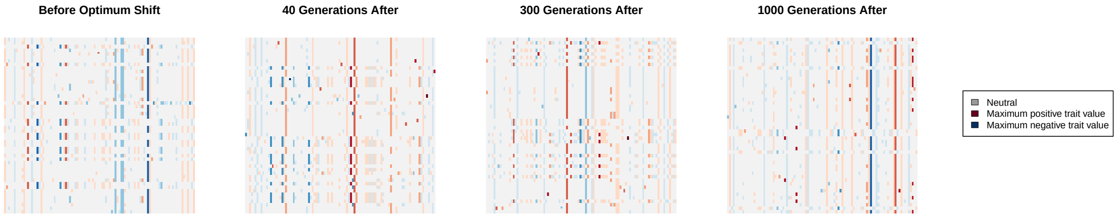


(b)

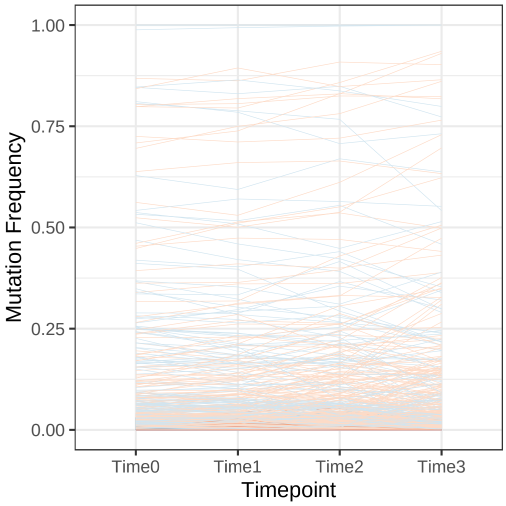


(c)

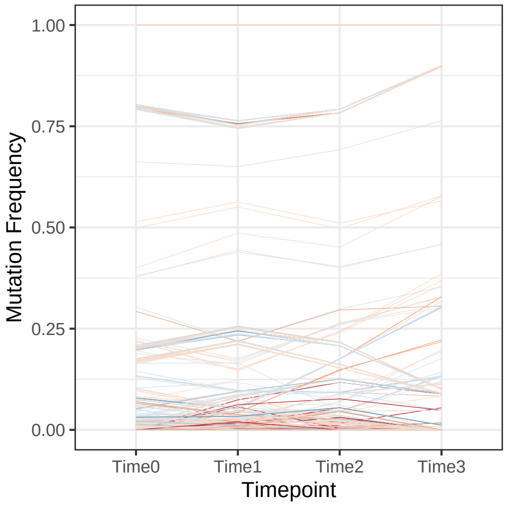


(d)

Figure F: (a), (b) Haplotype plots and (c), (d) allele–frequency changes for a population of size $N = 10,000$. Populations are either (a) outcrossing or (b) 99.9% self–fertilising.

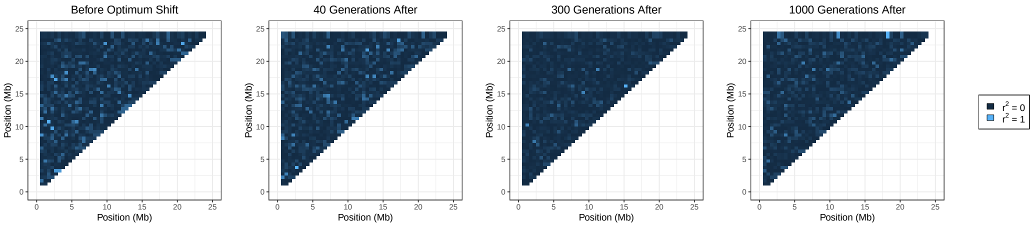


(a)

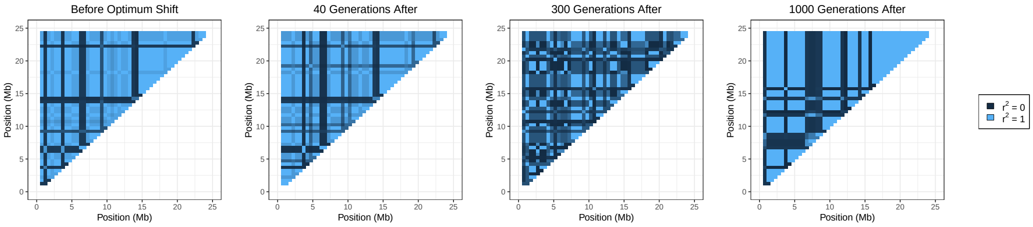


(b)

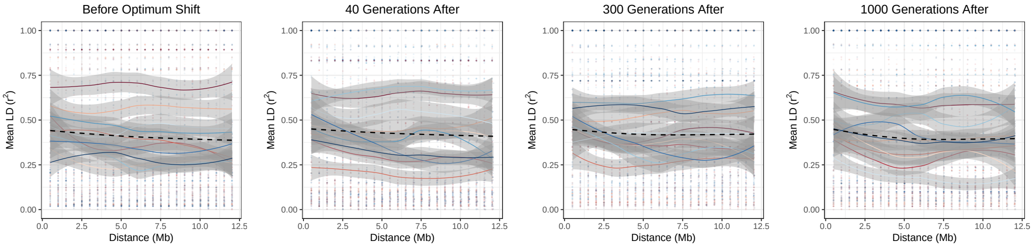


(c)

Figure G: (a), (b) LD heatmaps and (c) LD decay for a population of size $N = 10,000$. Populations are either (a) outcrossing or (b), (c) 99.9% self–fertilising.

### Additional figures with deleterious mutations present

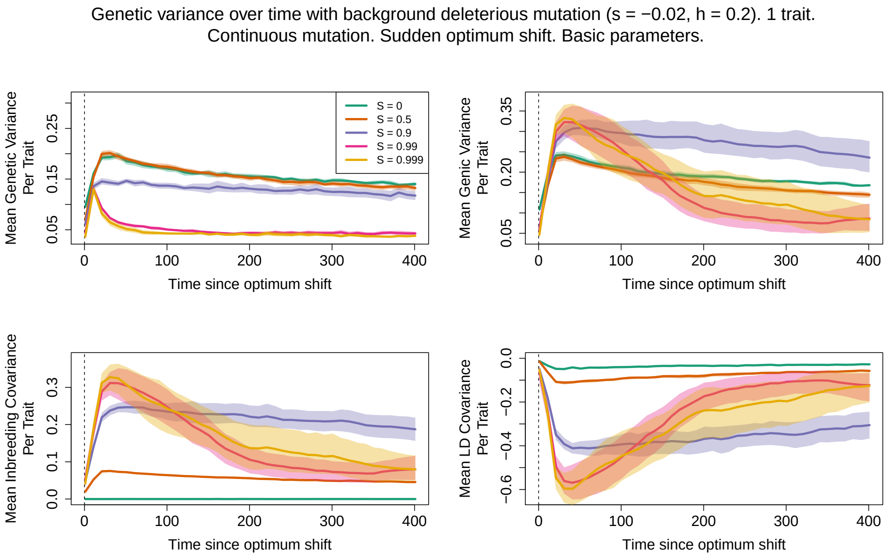


(a)

Figure H: Variance measurements when trait mutations are non–pleiotropic and background deleterious mutations are present with $h = 0.2$. Panel order is as in Figure 2 in the main text.

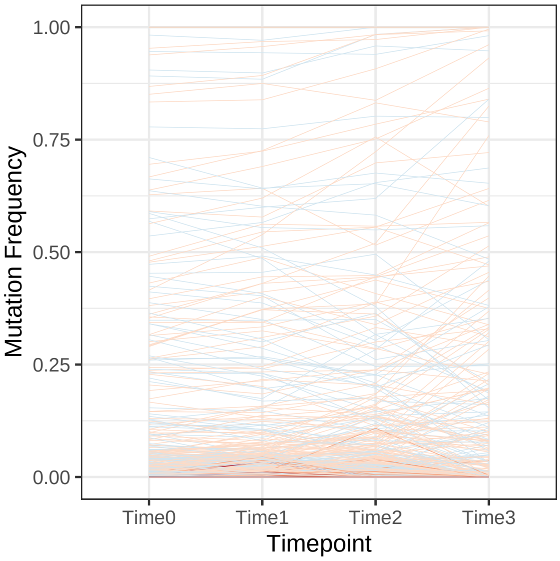


(a)

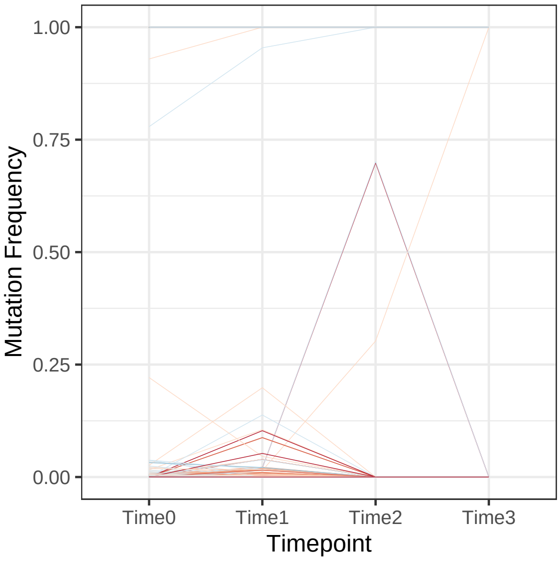


(b)

Figure I: Plots of change in trait mutation frequency over the timeframe of adaptation, when trait mutations are non–pleiotropic and background deleterious mutations are present with $h = 0.2$. The population is either (a) outcrossing, or (b) 99.9% self–fertilising.

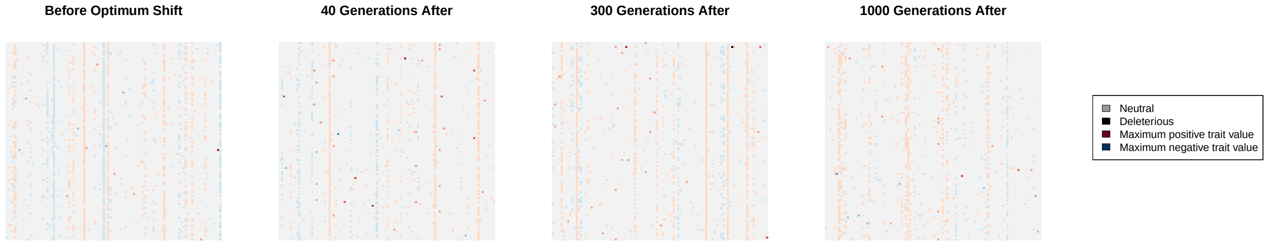


(a)

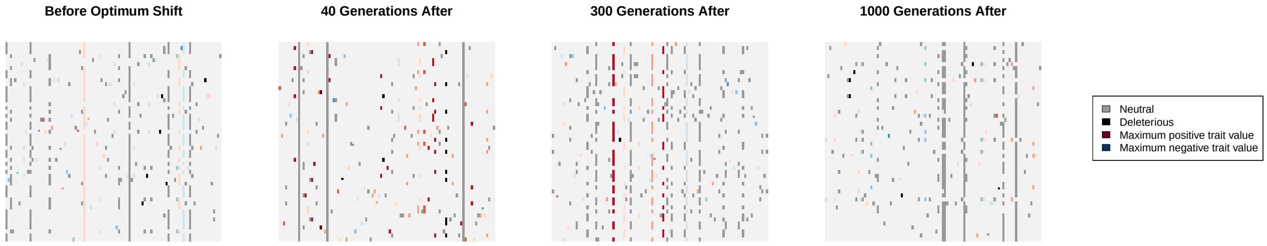


(b)

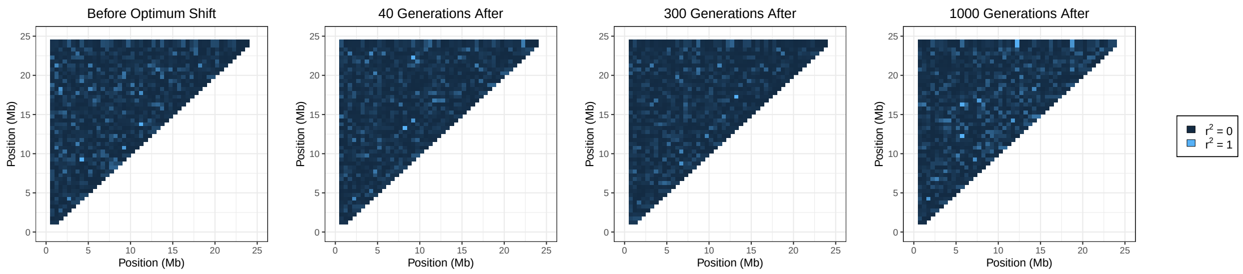


(c)

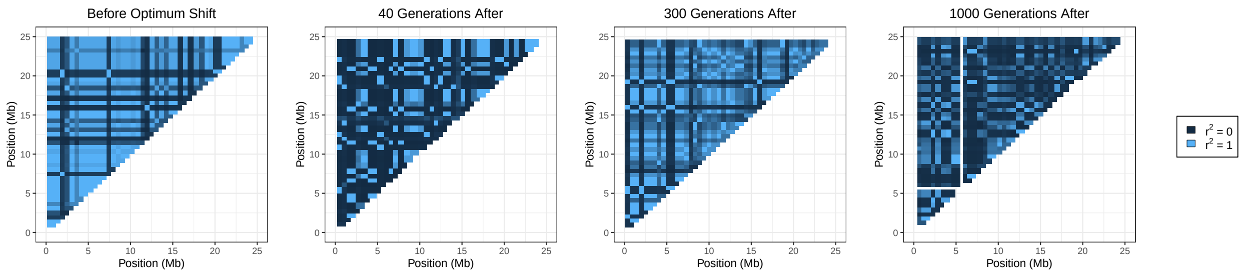


(d)

Figure J: (a), (b) Haplotype plots and (c), (d) LD heatmaps following a sudden change in optimum shift when trait mutations are pleiotropic and background deleterious mutations are present with $h = 0.02$. Populations are either (a), (c) outcrossing or (b), (d) 99.9% self–fertilising.

### Effect of rescaled parameters in outcrossing populations

#### Fitness and trait variance

Without background deleterious mutations, fitness measurements are generally similar between the selfing and rescaled outcrossing population (Figure K); the only visible contrast arise for variance in fitness under pleiotropy. Differences between cases become apparent when looking at variance measurements (Figure L). Genetic variance is slightly higher overall in the rescaled–outcrossing population, caused by a higher additive variance and LD covariance of a lower magnitude. Given that inbreeding variance is non–zero under the high selfing case, these results suggest that the inbreeding-based reduction in recombination causes a greater loss in trait variation than expected under simply reducing the recombination rate, reflecting purging of selected variants.

With recessive deleterious mutations included (Figure M), genic and genetic variance is higher under outcrossing, but LD covariance is more strongly negative. We also observe that the mean fitness is greatly reduced in the outcrossing case, and inbreeding depression has increased. Overall, this result suggests that low–recombination outcrossers more strongly suffer from selection interference, so recessive deleterious mutations persist leading to reduced fitness. This interference also means that trait variants with compensatory effects are more likely to be found on the same haplotypes. Conversely, these recessive deleterious mutations are purged under high self–fertilisation.

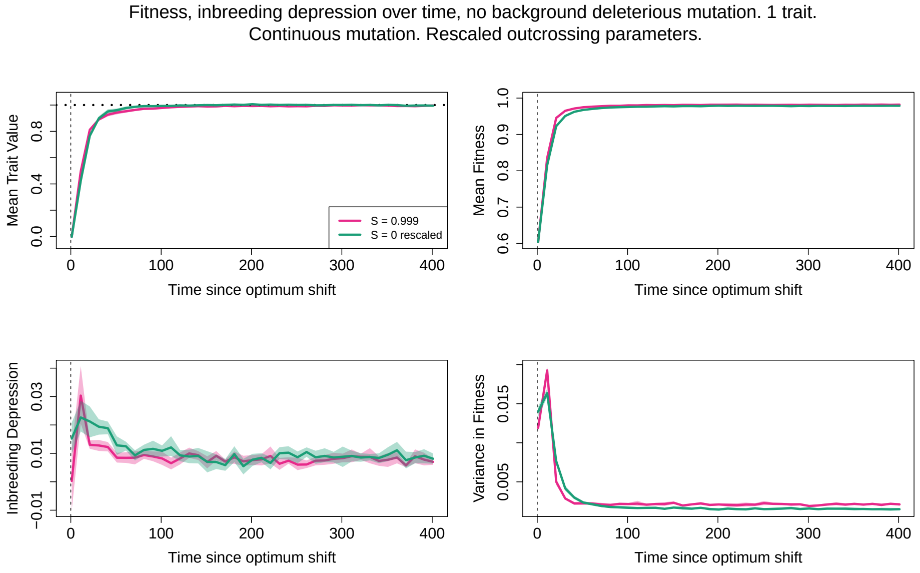


(a)

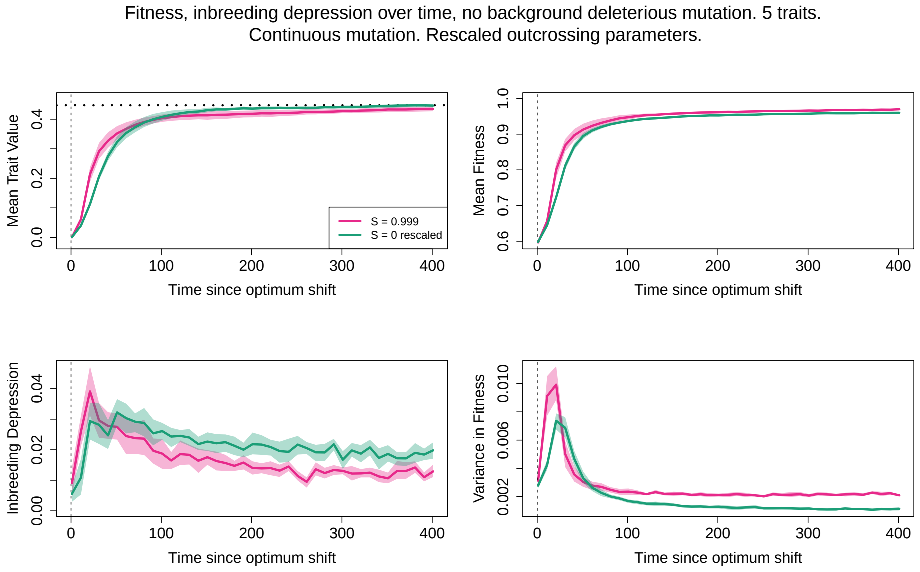


(b)

Figure K: As Figure 1 in the main text, but comparing the $S = 0.999$ case to an outcrossing population with rescaled mutation, recombination rates.

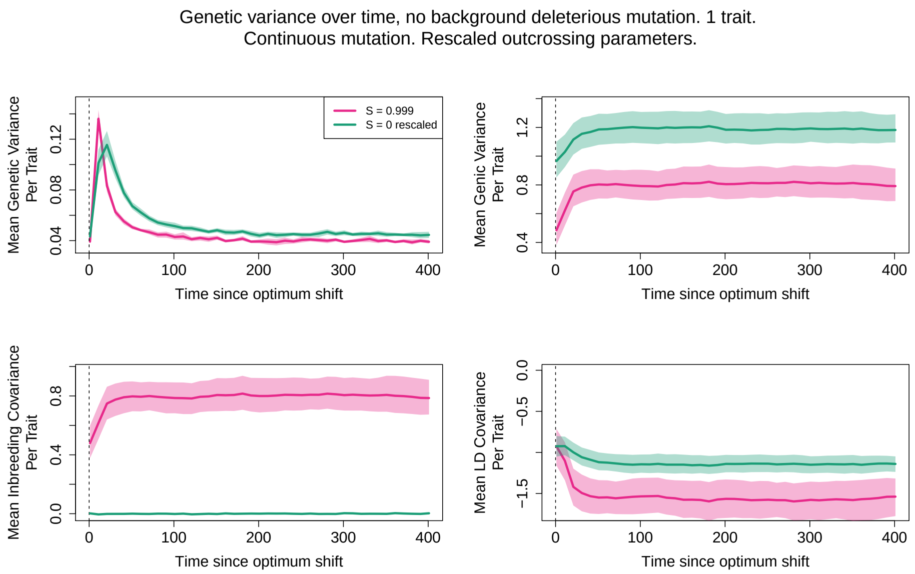


(a)

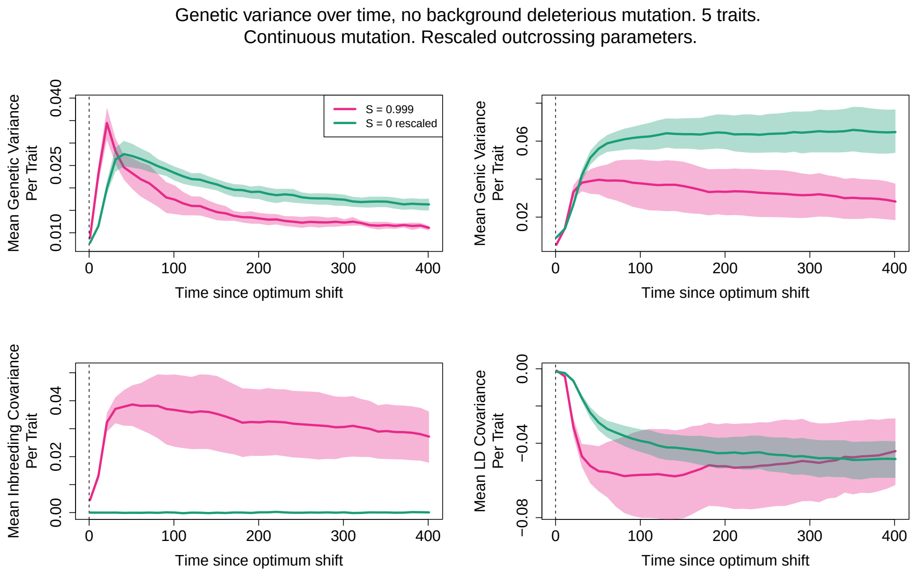


(b)

Figure L: Comparing variance components under very high self–fertilisation (99.9%) to an outcrossing population with rescaled mutation and recombination parameters. Deleterious mutations are absent. Pleiotropy is either (a) absent or (b) present.

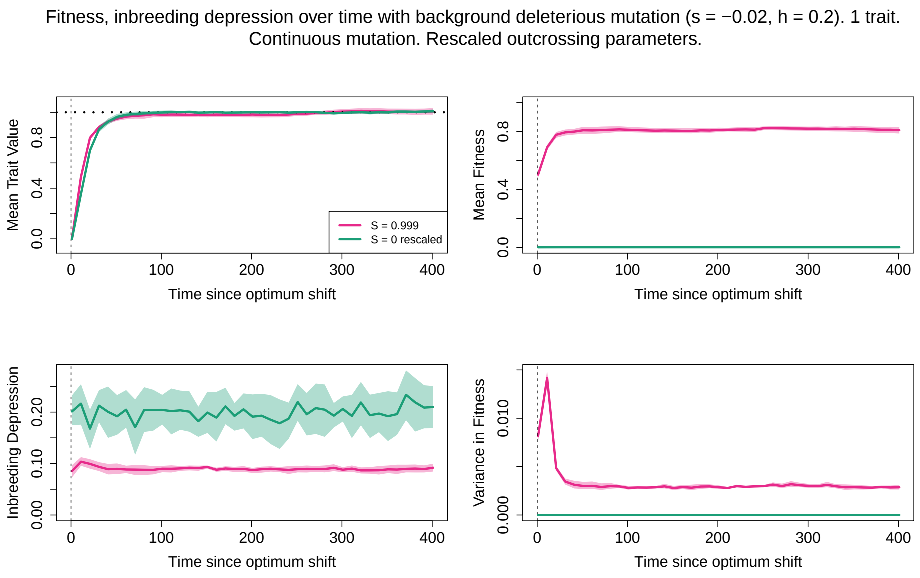


(a)

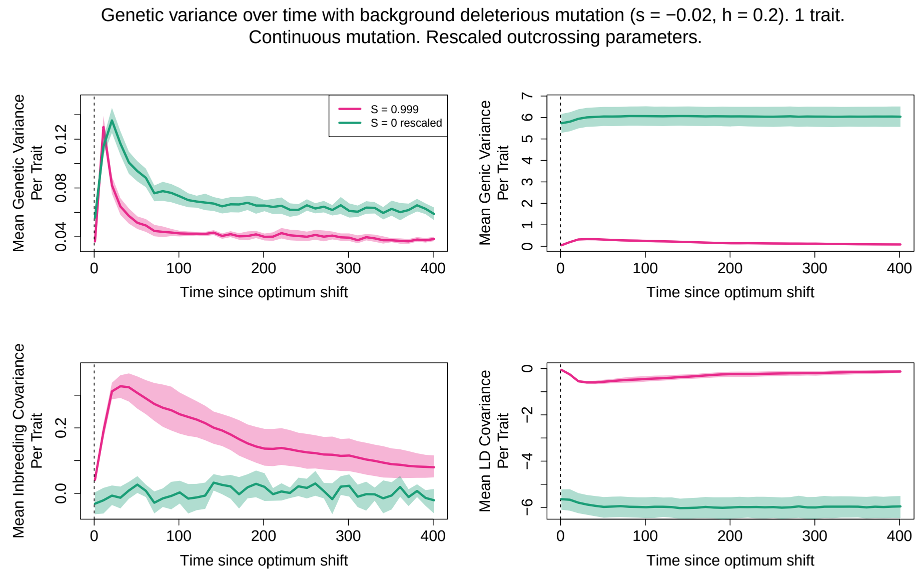


(b)

Figure M: (a) Fitness measurements (mean trait value; mean fitness; inbreeding depression; variance in fitness) and (b) genetic variance components for the rescaled outcrossing population, compared to the high self–fertilising population.

#### Haplotype structure

By inspecting the LD decay under the rescaled outcrossing case, we obtain further evidence that LD clustering does not occur to the same degree as under the high selfing case. In particular, when deleterious mutations are absent, then we do not observe persistence of LD over the same large distances as under high selfing (Figure N(a), N(b)), although some increased LD is formed over shorter distances as adaptation is ongoing, when pleiotropy is present. When recessive deleterious mutations are included in the non–pleiotropic case, we observe quite complex LD patterns; overall LD can be elevated quite significantly (e.g., at 300 generations after optimum shift), but is more likely to decay over longer distances, in contrast to the full selfing case (Figure N(c)). These results demonstrate that while some aspect of the high–LD behaviour can be formed in outcrossers with low recombination if background deleterious mutations are present, the exact formation of linkage–blocks in selfers is also affected by increased homozygosity due to inbreeding. Figures O, P show example haplotype plots and LD heatmaps for individual simulations.

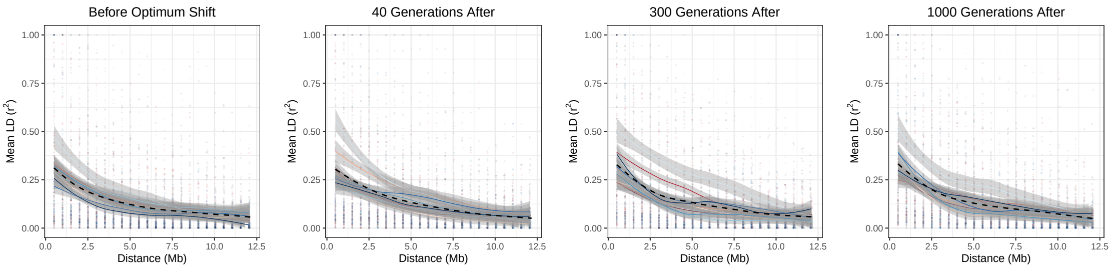


(a)

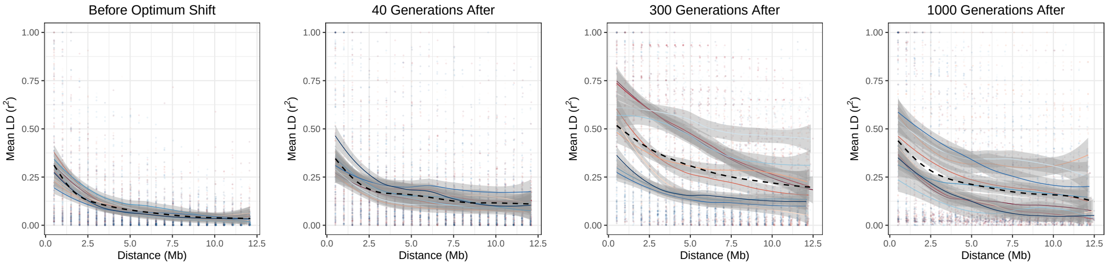


(b)

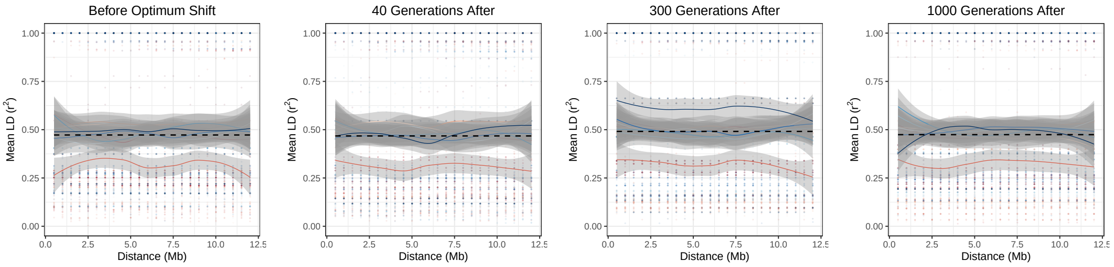


(c)

Figure N: LD decay for outcrossing populations with rescaled recombination, mutation rates to match those in a highly selfing population. Cases considered are (a) no deleterious mutations or pleiotropy; (b) no deleterious mutations, pleiotropy present; or (c) deleterious mutations (with $h = 0.2$) and pleiotropy absent.

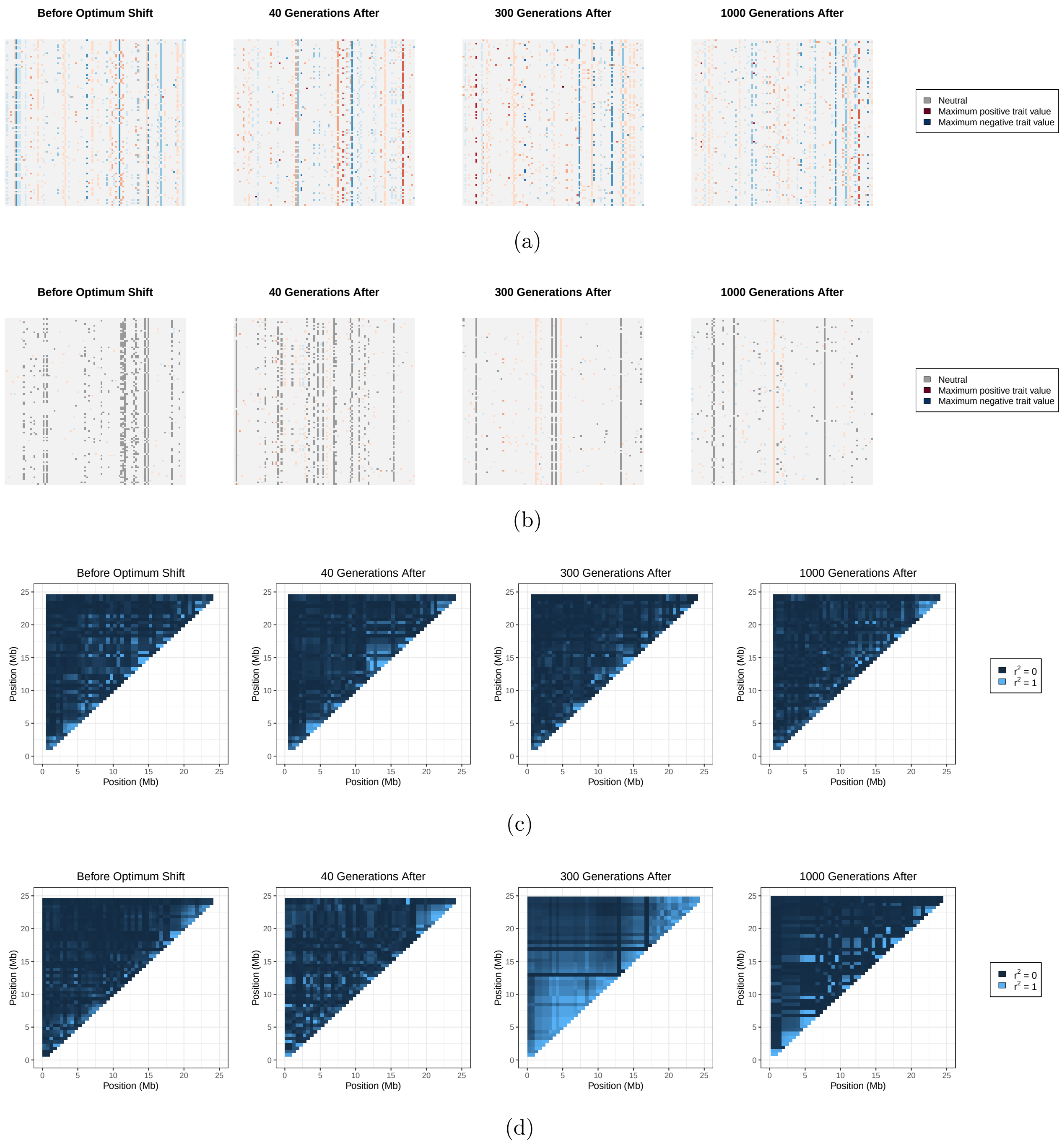


Figure O: Haplotype plots (a), (b) and LD heatmaps (c), (d) for outcrossing populations with rescaled recombination, mutation rates to match those in a highly selfing population. There are no background deleterious mutations, and pleiotropy is either absent (a) (c) or present (b) (d).

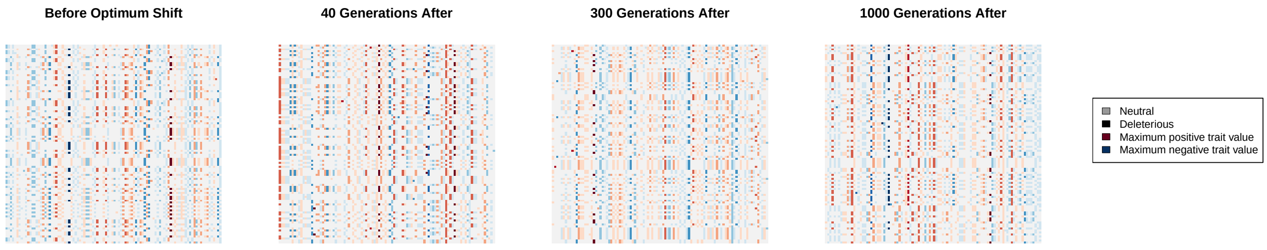


(a)

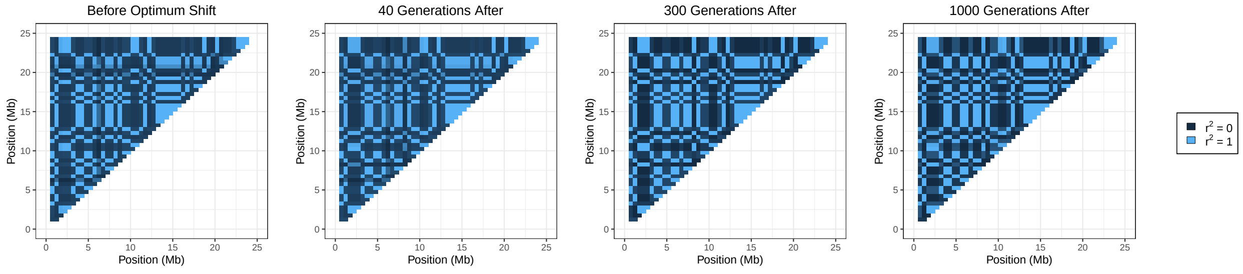


(b)

Figure P: As Fig O but only for the non–pleiotropic case and with deleterious mutations included with $h = 0.2$.

### Additional method figures

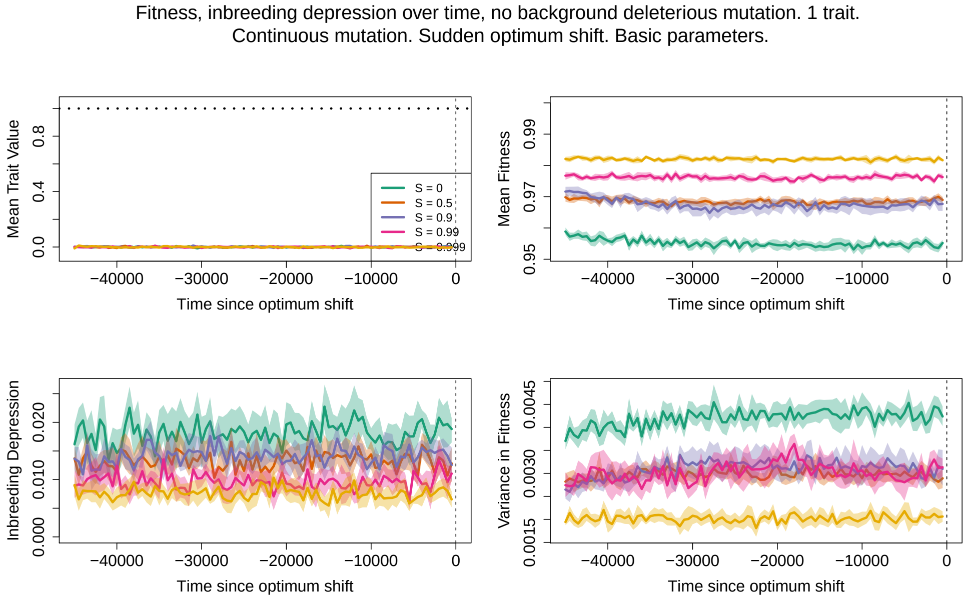


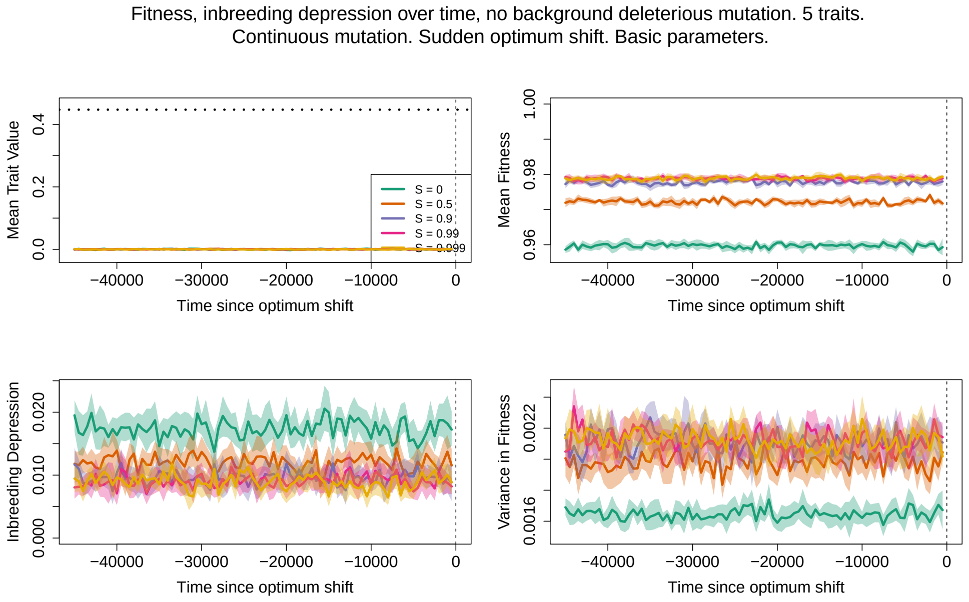


Figure Q: As Figure 1 in the main text, but instead tracking fitness measurements before the optimum shift.

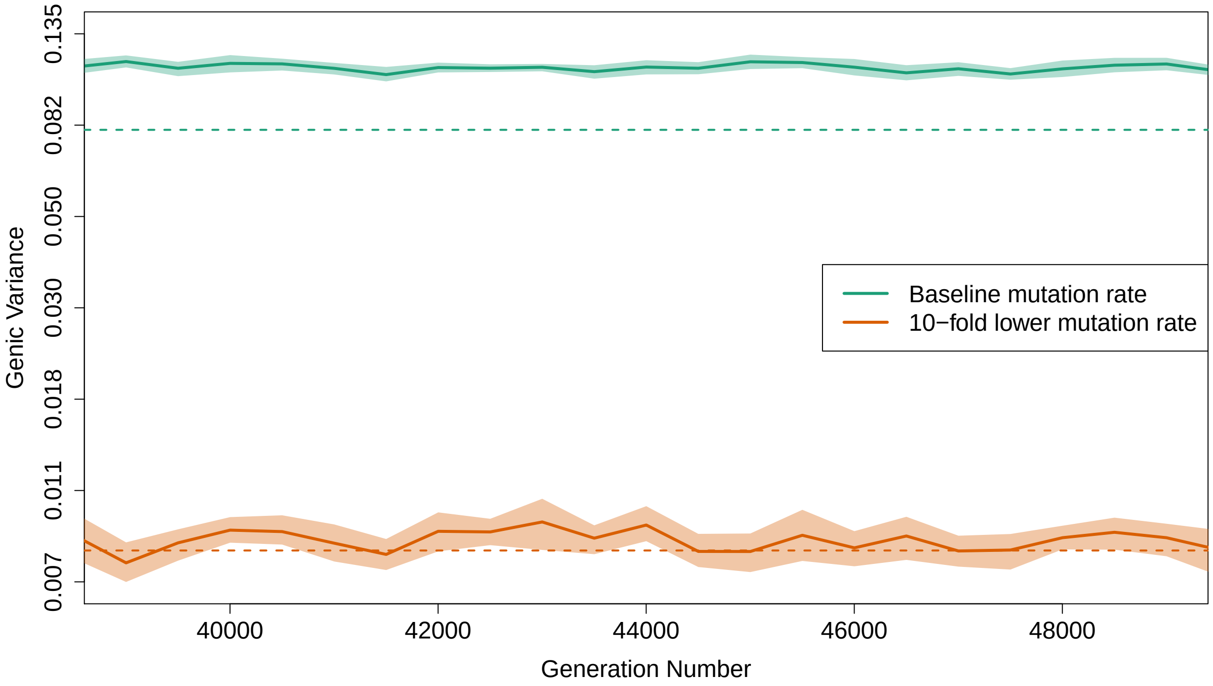


Figure R: Comparing genic variance just before the optimum shift, averaged over 10 simulation replicates (solid lines, bands show 95% confidence intervals) to the house–of–cards expectation (dotted lines). There is no pleiotropy; populations are outcrossing; and no background deleterious mutations are present. Simulations either use the standard mutation and recombination rates (green) or 10–fold lower values (orange). Note the Y–axis is on a natural–log scale.
